# Understanding of ATP-lid conformational dynamics in the N-terminal domain of Hsp90 and its mutants by use of a computational biochemistry approach

**DOI:** 10.1101/2025.02.12.637981

**Authors:** Keigo Gohda

## Abstract

Chaperone Hsp90 regulates the activation and maturation of various protein, and is an attractive target for drug discovery. In catalytic cycle of Hsp90, ATP hydrolysis is a key event that drives structural changes, including the interchange of the dimeric Hsp90 structure between open and closed forms. For ATP hydrolysis, ATP-lid closure in the ATP-binding site from the up- to down-conformation is an indispensable co-ordinated structural change. However, the atomistic mechanism underlying lid closure remains unclear. In this study, a computational biochemistry approach was applied to wild-type apo and ATP-complex structures, and the lid-mutants A107N and T101I ATP-complex, to understand lid closure.

A total of 15-μs molecular dynamic simulation, including the equilibration and production phases, was conducted for every structure starting from the lid up-conformation, but no lid-closures were observed. However, a very early event, i.e., *a sign*, of lid closure have been captured. In the simulations of wild-type and A107N ATP-complex structures, lid segment showed conformational fluctuations, and helix-7 (H7) segment in lid segment was unwound. This conformational instability of lid segment energetically weakened its interaction with facing region, suggesting the up-to-down transition was triggered by the instability of lid segment, particularly H7 segment. The interaction energy between lid segment and facing region was ranked in the order A107N > wild-type > T101I, which was correlated with experimental results such as the orders of ATP-hydrolytic activity and rapidity of conformational changes of lid segment, measured in ATP-spiking experiments by fluorescence resonance energy transfer method (i.e., A107N > wild-type > T101I).

## 1. Introduction

The 90 kDa heat shock protein (Hsp90) is a chaperone protein that regulates the activation and maturation of various proteins, i.e., clients (Biebl & Buchner, 2019; Prodromou, 2016). Because client proteins play central roles in essential cellular processes involved in cell growth and proliferation, Hsp90 is an attractive target for drug discovery, especially in cancer and neurodegeneration (Hoy, 2022; Wei at al., 2024; Xiao & Liu, 2020). Hsp90 uses ATP molecules in its chaperone function and is composed of an N-terminal domain (NTD), including the ATP-binding site; a middle domain that mainly contributes to interactions with clients and co-chaperones; and a C-terminal domain (CTD) (Meyer et al., 2003; Nemoto et al., 1995; Prodromou et al., 1997a). Hsp90 functions as a homodimer, and the CTD is primarily involved in the dimerization process, forming one end of a molecular clamp (Ali et al., 2006; Prodromou et al., 2000).

In apo state, i.e., no ATP molecules bound to the ATP-binding site, the dimeric Hsp90 subunits adopt “open form” (Prodromou & Bjorklund, 2022). Binding an ATP molecule to the NTD drives conformational changes of the dimeric structure to “closed form”, including NTD dimerization, which forms the other end of the molecular clamp (Ali et al., 2006). The closed form activates the catalytic site, leading to hydrolysis of ATP molecules and completion of the catalytic cycle, that is, the release of ADP molecules and mono-phosphates from the dimeric subunits. Various cochaperone proteins are involved in the catalytic cycle (Li & Buchner, 2013). The rate-limiting step of the cycle involves co-ordinated structural changes during the opening–closing process. The co-ordinated structural changes involve a closure of lid segment (Gly94-Gly123) in the NTD, which covers over a bound ATP molecule in the ATP-binding site, a swap of N-terminal β-strands crossing between NTD monomers, and an association between the NTD and middle domain **(Figure 1)**. These structural changes are collectively slow and interdependent (Schubert et al., 2021; Schulze et al., 2016) and complete a proper placement of catalytic residues such as Arg32, Glu33, and Arg380 (Mader et al., 2020; Prodromou et al., 1997a; Prodromou & Bjorklund, 2022) (yeast Hsp90 numbering is used throughout this manuscript).

**Figure 1.**
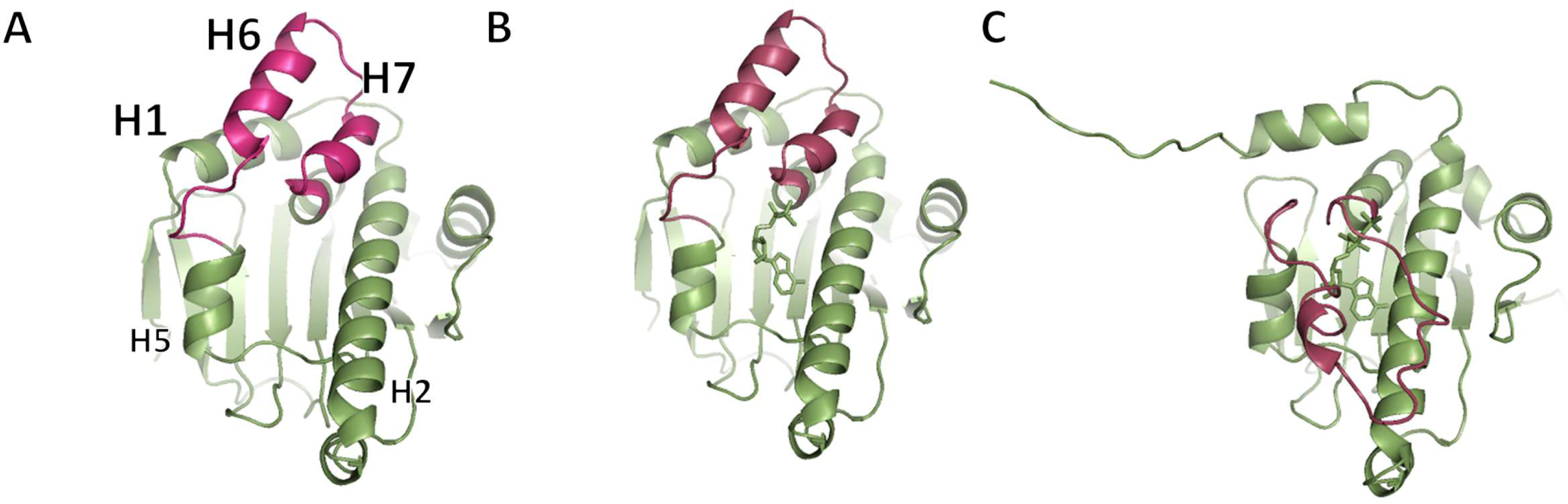
The X-ray structures of Hsp90 NTD. Lid segment (Gly94-Gly123) is colored in red. (A) apo structure with “up” conformation of lid segment (1AH6; Prodromou et al., 1997b), (B) ADP-complex structure with “up” conformation (1AM1; Prodromou et al., 1997a), and (C) ATP-complex structure with “down” conformation, which is extracted from the X-ray dimeric Hsp90/Sba1 complex structure (2CG9; Ali et al., 2006).

However, the atomistic mechanism of the co-ordinated structural changes has not yet been elucidated. Among these structural changes, we focused on understanding the ATP lid closure, which is indispensable for ATP hydrolysis. In X-ray and cryo-electron microscopic structures of Hsp90 monomeric or dimeric subunit, the ATP-lid segment takes open conformation in the open form of the dimeric subunit and closed conformation in the closed dimeric subunit (Biebl & Buchner, 2019; Prodromou & Bjorklund, 2022) (In this manuscript, the open and closed conformations of the ATP-lid segment are named as "up" and "down" conformations to avoid confusion with structural change of the dimeric subunit). In human Hsp90, it was recently reported, based on NMR measurements, that the up-and-down conformational transition of the ATP-lid occurs in the apo state of the NTD (Henot et al., 2022). However, no information is available on the transition process in the ATP-complex structure. As the ATP binding triggers the co-ordinated structural changes necessary for ATP hydrolysis, an ATP molecule may induce a lid conformational transition in the up-conformation. To understand the transition mechanism at atomistic scale, 15-μs length conventional molecular dynamic (MD) simulation, including the equilibration and production phases, was implemented for apo and ATP-complex structures of a monomeric NTD of yeast Hsp90.

Mutants in the lid segment that affect the ATP hydrolytic activity are known among the identified Hsp90 mutants (Nathan & Lindquist, 1995; Prodromou et al., 2000; Reidy & Masison, 2020; Richter et al., 2022; Siligardi et al., 2004). T101I and A107N mutants modulate ATP hydrolytic activity in the direction of attenuation and enhancement, respectively (Prodromou et al., 2000). However, there is no structural information on these mutants, and it is unclear how mutations in the lid segment affect ATP hydrolytic activity at the atomistic level. Since elucidation of the modulating mechanism of the lid mutants may contribute to a comprehensive understanding of lid function, MD simulations were performed for the modeled ATP-complex structures of T101I and A107N.

## 2. Materials and methods

### 2.1. Materials

The X-ray structures of yeast Hsp90 apo and ADP-complex NTDs were used in this study (PDB IDs: 1AH6 and 1AM1) (Prodromou et al., 1997a; Prodromou et al., 1997b). To prepare the structures for MD simulation, nucleotides, water, and ions were removed from each X-ray structure, and hydrogen atoms were added. The protonation states of the side-chain atoms were assessed using the PDB2PQR server (Dolinsky et al., 2007). In assigning the protonation states, a neutral pH of 7.0 was assumed. *In silico* mutant structures were modeled by altering the amino acids at relevant positions in the X-ray structures. The model structures were manipulated using the MAESTRO modeling package (Schrodinger LLC, New York, NY, 2020). All molecular mechanics calculations were conducted using the *pmemd*.*cuda* or *sander* modules of AMBER22/AMBERTOOL22 (Case et al., 2022; Goetz et al., 2012; Le Grand et al., 2013; Salomon-Ferrer et al., 2013).

For computations using AMBER22, the *tleap* module was used to prepare the topology and coordinate files of the proteins and ligands. The ff14SB force field and TIP3P model (Jorgensen et al., 1983; Maier et al., 2015) were used for protein and water molecules, respectively. The topology and parameter files of the ATP molecules and a Mg^2+^ ion were obtained from AMBER parameter database of Bryce’s group at the University of Manchester (Alln et al., 2012; Bryce, 2008; Meagher et al., 2003). The initial coordinates of the phosphate group in ATP molecule were adapted from human Hsp90α X-ray NTD structure complexed with ATP (PDB ID: 3T0Z) (Li et al., 2012).

The model structures prepared using the *tleap* module were immersed in an octahedral box containing water molecules to impose a periodic boundary. The largest distance between the complex and the water molecules in the box was set to 12.0 Å. To neutralize the entire system, Na^+^ counterions were added to the box (Ibragimova & Wade, 1998).

### 2.2. MD Simulation

The model structure in an explicit solvent was subjected to energy minimization. The minimization procedure was divided into the following three steps with/without constraints: (a) the positions of all heavy atoms in the protein and ATP were fixed (positionally constrained with a 50 kcal/mol·Å^2^ weight), and hydrogen atoms, ionic atoms, and water molecules were allowed to move freely, (b) weak constraints were added to all heavy atoms in the protein and ATP (positionally constrained with a 2 kcal/mol·Å^2^ weight), and (c) no constraints were added. Each minimization process employed 500 cycles of the steepest-descent method, followed by the conjugate-gradient method, until 3000 cycles were completed. The energy convergence criterion gradient in each minimization step was 0.0001 kcal/mol·Å for the root-mean-square of the Cartesian elements of the gradient. The cutoff distance for nonbonding interactions was 8.0 Å, and the nonbonding interaction list was updated every 25 steps.

The molecular dynamics calculations for the minimized structure were conducted in three consecutive steps: heating, density, and simulations. The time step in the MD simulation was 2 fs, and the SHAKE constraint was used for bonds involving hydrogen atoms (Ryckaert et al., 1977). The particle-mesh Ewald method was used to calculate electrostatic energy (Darden et al., 1993; Essmann et al., 1995). The system was heated from 0 to 300 K over 100 ps using the Andersen temperature-coupling system (Andrea et al., 1983). The velocities were randomized every 2 ps for temperature scaling. The Cα atoms of the main-chain atoms in a protein were positionally constrained by a 2 kcal/mol·Å^2^ weight. After heating, the system was equilibrated at 300 K at 1 bar for 50 ps. The constant-pressure periodic boundary condition was applied using a Berendsen barostat with a 1 ps pressure relaxation (Berendsen et al., 1984). The same positional constraints were applied to the system as the heating step. Finally, a 50 ns simulation without any constraints was performed, and the obtained structures were applied for further MD simulations.

All statistical analyses were performed using the Student’s t-test with MedCalc22.020 (MedCalc Software Ltd., Belgium, 2024).

### 2.3. Helicity assessment

The degree of helical contents of helix segments was defined as ‘helicity’. A helicity by residue was assessed as a ‘residual helicity’ as to whether the conformation of a residue took a helical profile. The helical profile was estimated using two structural parameters derived from Kabsch and Sander’s secondary structural propensity: hydrogen-bonded and geometrical structures (Kabsch & Sander, 1983).

First, according to the Kabsch and Sander’s propensity, a hydrogen-bonded structure is defined as a structure that adopts internal hydrogen bonds within five consecutive residues, assessed by the formation of hydrogen bonds between the backbone amide (N−H) of the *i*th residue and the carbonyl (C=O) atom of the *j*th residue (*j* = *i* + 3, 4, or 5). Residue secondary-structural types in the Kabsch and Sander’s propensity were assigned as 3−10 helix (G), α-helix (H), π-helix (I), turn (T), parallel β-sheet (E), antiparallel β-sheet (B), bend (S), or no structure. In this study, if the hydrogen-bonded structure of a residue (*i*) was assessed as G, H, I, or T, this residue was assigned as a ‘helical hydrogen-bonded’ residue.

Second, a geometrical structure was assessed by the chirality of main-chain structure as to whether a right- or left-handed conformation was present by assessing a dihedral angle composed of four consecutive Cα atoms ([*i* − 1]th to [*i* + 2]th) (Kabsch & Sander, 1983). In this study, if the geometrical structure of a residue (*i*) was a positive dihedral angle, the residue was assigned as a ‘right-handed helical’ residue.

Using the two residual structural assignments, the residual helicity of a residue was defined as “plus one” (+1) if the residue was assigned to be ‘helical hydrogen-bonded’ and ‘right-handed helical’ in both residual assignments. Subsequently, every residual helicity of the entire helical segment was combined to provide the total helicity of the helical segment.

In this study, the complete scores of the total helicity for the helix-1 (H1), helix-6 (H6) and helix-7 (H7) segments were 12, 12, and 6 because each segment was composed of 12 residues (Ala10 to Asn21), 12 residues (Ser99 to Ala110), and 6 residues (Val114 to Phe120), respectively. The helicity ratio at each time point was then calculated by dividing the helicity score of a segment by the complete score of the segment.

### 2.4. Principal component analysis

Principal component analysis (PCA) was calculated using *cpptraj* module (Roe & Cheatham, 2013) and visualized using *colormap.py* of AMBER22. The principal modes were calculated by diagonalizing the covariance matrix of the fluctuations obtained from the atomic coordinates to produce the eigenvectors.

### 2.5. MMGBSA calculation

The molecular mechanics/generalized Born surface area (MMGBSA) method was used to estimate the interaction energy (Gohlke et al., 2003; Gohlke & Case, 2004). An MMGBSA calculation was conducted using the *MMPBSA.py* module of AMBER22 (Miller et al., 2012) for every 100 ps-snapshot in the MD trajectory. For the generalized Born term, the Hawkins, Cramer, and Truhlar pairwise generalized Born model was used (Hawkins et al., 1995 Hawkins et al., 1996), with the parameters described by Tsui and Case (Tsui & Case, 2001). The salt concentration in the solution was set to be 0.1 M.

The MMGBSA energy interaction (dG_inter_) between the lid segment and facing region was calculated using the following formula:

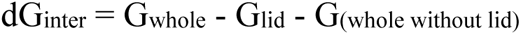

## 3. Results

### 3.1. Effect of ATP binding on NTD structure

#### 3.1.1. Preparation of apo and ATP-complex structures for MD simulation

For apo structure, the X-ray structure of yeast apo NTD (PDB ID: 1AH6) (Prodromou et al., 1997b) was used and subjected to 5 μs MD simulation as an equilibration phase. During equilibration, the entire structure remained stable, and the degree of helical content (helicity) remained unchanged **(Figure 2 A,B,D)**. However, H6 (Ser99 to Ala110) and H7 (Val114 to Phe120) segments in the lid segment were displaced from the initial position of the X-ray structure since around 4 μs **(Figure 2 C)**. The helicity of H1 (Ala10 to Asn21) and lid segments (H6 and H7 segments) was quantified using the helicity assessment method (Gohda, 2022) and was stable during equilibration. Helicity assessment was performed for two helical structural parameters based on Kabsch and Sander’s secondary structural propensity: hydrogen-bonded and geometrical structures (Kabsch & Sander, 1983) (see Materials and Methods).

**Figure 2.**
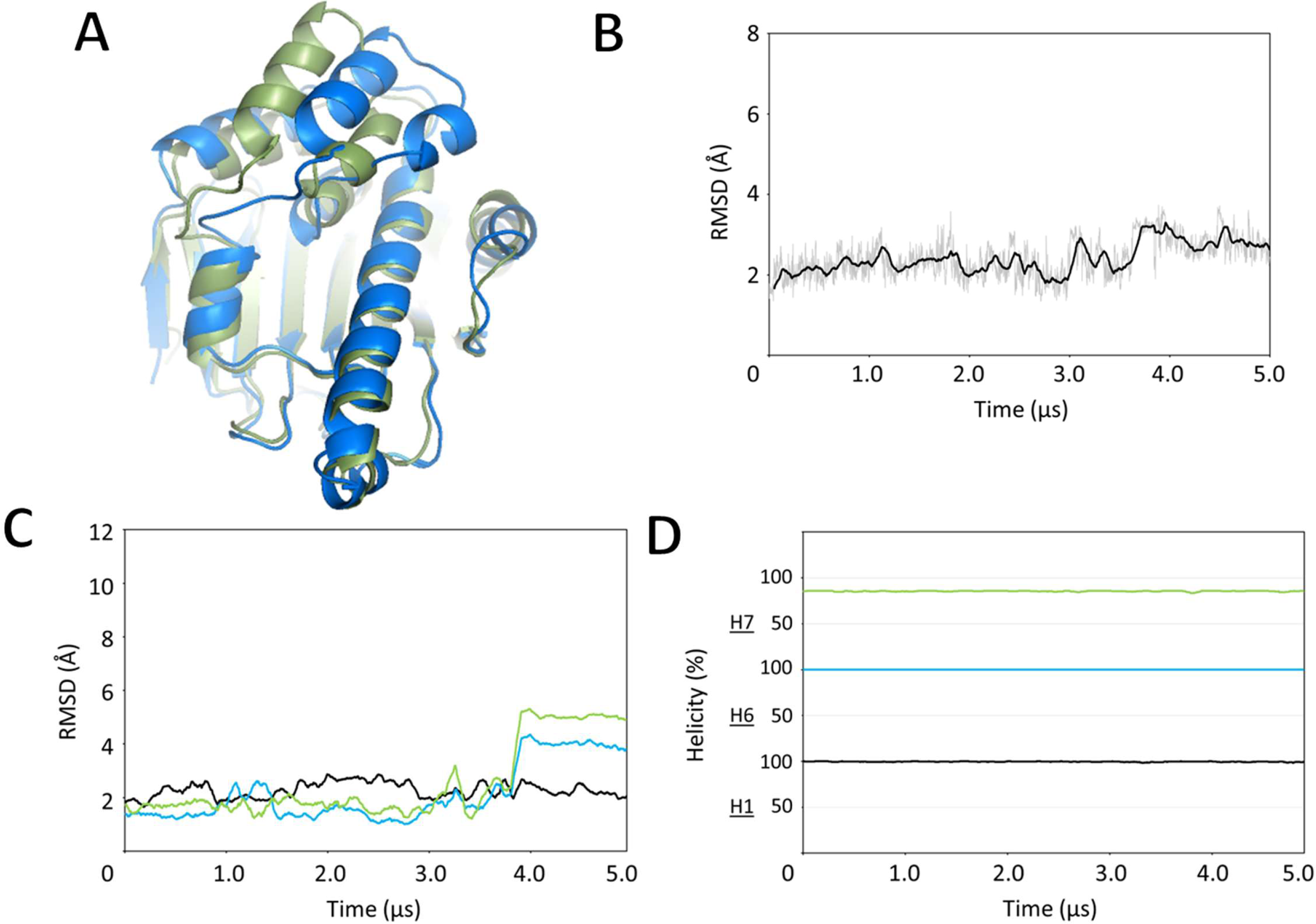
MD equilibration for the apo structure of the NTD. (A) apo structure after equilibration (blue) compared with the initial apo structure (green), the RMSD trajectories of the entire structure (B) and the H1/H6/H7 segments (C), and (D) the helicity trajectories of the H1/H6/H7 segments. Plots of the H1/H6/H7 segments are colored in black, light blue and light green, respectively.

Visual inspection of the initial X-ray apo structure showed that the symmetry-related NTD structures were located close to the lid segment of the parent structure **(Figure S1)**. Thus, the equilibrated structure of apo NTD at 5 μs was considered to be a free structure from the crystal packing. A similar observation of NTD structure crystal packing was previously reported for the H6 conformation (Stachowski & Fischer, 2022). The equilibrated apo structure was used for the production phase of the MD simulations.

For the ATP-complex, the X-ray structure of the yeast ADP-complex NTD (PDB ID: 1AM1) (Prodromou et al., 1997a) was used instead of the ATP-complex structure, which has not yet been determined. The positions of γ-phosphate, a magnesium ion, and two coordinating water molecules for the ATP-complex were adapted from human Hsp90α X-ray NTD structure complexed with ATP (PDB ID: 3T0Z) (Li et al., 2012), and information from a previous study on conformations of the catalytic residues was used for model building (Mader et al., 2020; Prodromou & Bjorklund, 2022). The modeled ATP-complex structure was then subjected to 5 μs MD simulation for an equilibration phase. During equilibration, the root-mean-squared deviation (RMSD) of the entire structure was stable, but the helicity and position of the H7 segment changed and displaced (Figure 3).

**Figure 3.**
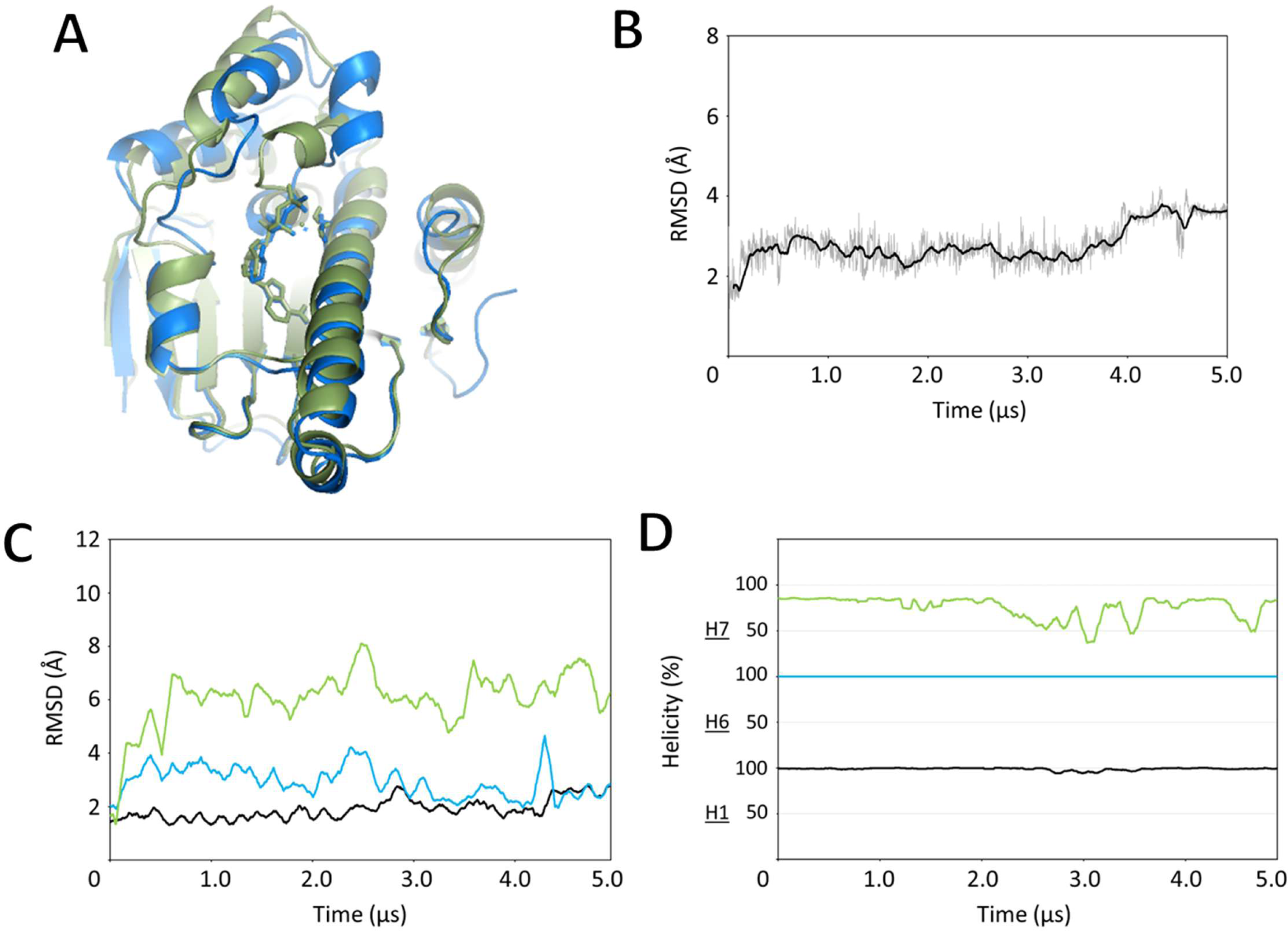
MD equilibration for the ATP-complex structure of the NTD. (A) ATP-complex structure after equilibration (blue) compared with the initial ATP-complex structure (green), the RMSD trajectories of the entire structure (B) and the H1/H6/H7 segments (C), and (D) the helicity trajectories of the H1/H6/H7 segments. Plots of the H1/H6/H7 segments are colored in black, light blue and light green, respectively.

Displacement of the lid segment due to the crystal packing appeared to be less than that observed in the apo structure because of the stability of the H6 segment. This may be due to the presence of ATP molecules at the ATP-binding site. Similar to the apo structure, the equilibrated ATP-complex structure was used for the production phase of the MD simulation.

#### 3.1.2. 10-μs production simulation of apo and ATP-complex structures

To investigate the effect of ATP binding on NTD structures, a long 10-μs MD simulation was conducted for the apo and ATP-complex structures. Production trajectories were analyzed in terms of conformational and energetic changes in the entire structure or in the local lid-conformation.

During the production of the apo and ATP-complex structures, both structures were stable, and the transition of the lid segment from the up- to down-conformation was not observed **(Figure 4 A,B)**. The RMSD values of the entire structure to each equilibrated structures were 2.21±0.470 Å and 2.73±0.792 Å in the apo and ATP-complex, respectively **(Figure S2)**. The RMSD of the lid segment in the ATP-complex was slightly larger than that in the apo structure (1.21±0.503 Å and 2.77±0.658 Å in the apo and ATP-complex structures, respectively). In particular, the H7 segment in the ATP-complex structure was unwound after 8 μs simulation, with the largest RMSD value (3.04±1.037 Å) among H1/H6/H7 segments of the apo and ATP-complex structures. By comparing a 10-μs snapshot in the production trajectory between the ATP-complex and apo structures, it was observed that the H7 segment in the ATP-complex was slightly moved away from the facing region to lid segment **(Figure 4 C)**. To clarify how the H7 segment leaves the facing region, the distance between the H7 segment and the facing region was examined. Two residues, Ile117 in the H7 segment and Ile29 in the H2 segment in the facing region, were selected as representative residues. They were placed in a face-to-face position **(Figure 4 C)**. During the production simulation, the average distances between Ile117 and Ile29 in the ATP-complex and apo structures were 10.32±1.744 Å and 8.45±0.650 Å, respectively.

**Figure 4.**
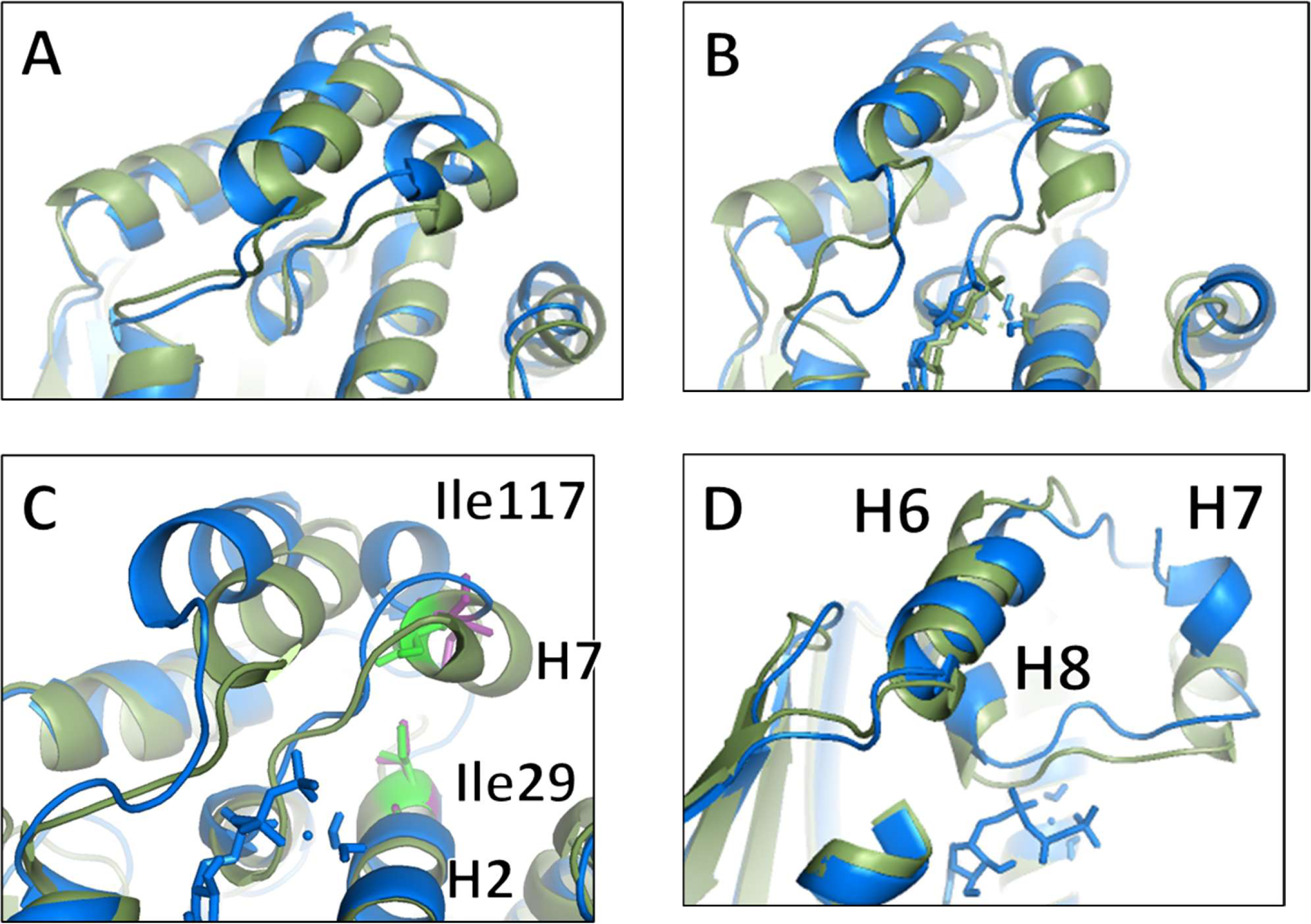
MD production for the apo and ATP-complex structures. (A) apo structure after production (blue) compared with the equilibrated apo structure (green), (B) ATP-complex structure after production (blue) compared with the equilibrated ATP-complex structure (green), (C) a close-up view of the H7 segment and facing region of the ATP-complex (blue) and apo structures (green). Ile117 on the H7 segment and Ile29 on the H2 segment in the ATP-complex structure were colored in purple, and those in the apo structure were colored in light green, and (D) a close-up view of a lid loop between the H7 and H8 segments in the ATP-complex structure after production (blue) overlapped to the apo structure after production (green).

In addition, the lid loop between segments H7 and H8 is pushed up in the ATP-complex structure in the direction of the H6 and H7 segments **(Figure 4 D)**. The side-chain of Lys98 in a lid loop-segment between segments H5 and H6 was found to ionically interact with the γ-phosphate group of the ATP molecule in the ATP-complex structure **(Figure S3)**. This interaction was continually observed during the production simulation. No other direct interaction was observed between the γ-phosphate and lid segment in the ATP-complex structure.

To understand the conformational landscape of the ATP-complex structure, PCA was implemented for the 10-μs trajectory. In the projection map, there were two large clusters, and the initial structure (0 μs) and final structure (10 μs) were located in each of the two clusters (**Figure 5 A**). As described above, H7 segment at 10 μs was unwound. This result thus suggested that unique structures in the simulation well identified in the map. Furthermore, the structures in the PCA projection were separately depicted for the first (0 μs to 5 μs) and late (6 μs to 10 μs) periods (**Figure 5 B**). The projection maps showed that the distribution of the structures was clearly different between the first and late periods. In the first period, most of the structures were populated in the cluster where the initial structure belonged. Contrarily, the structures in the late period were widely distributed in the map, indicating the conformational change in H7 segment mainly occurred in the late period of the simulation.

**Figure 5.**
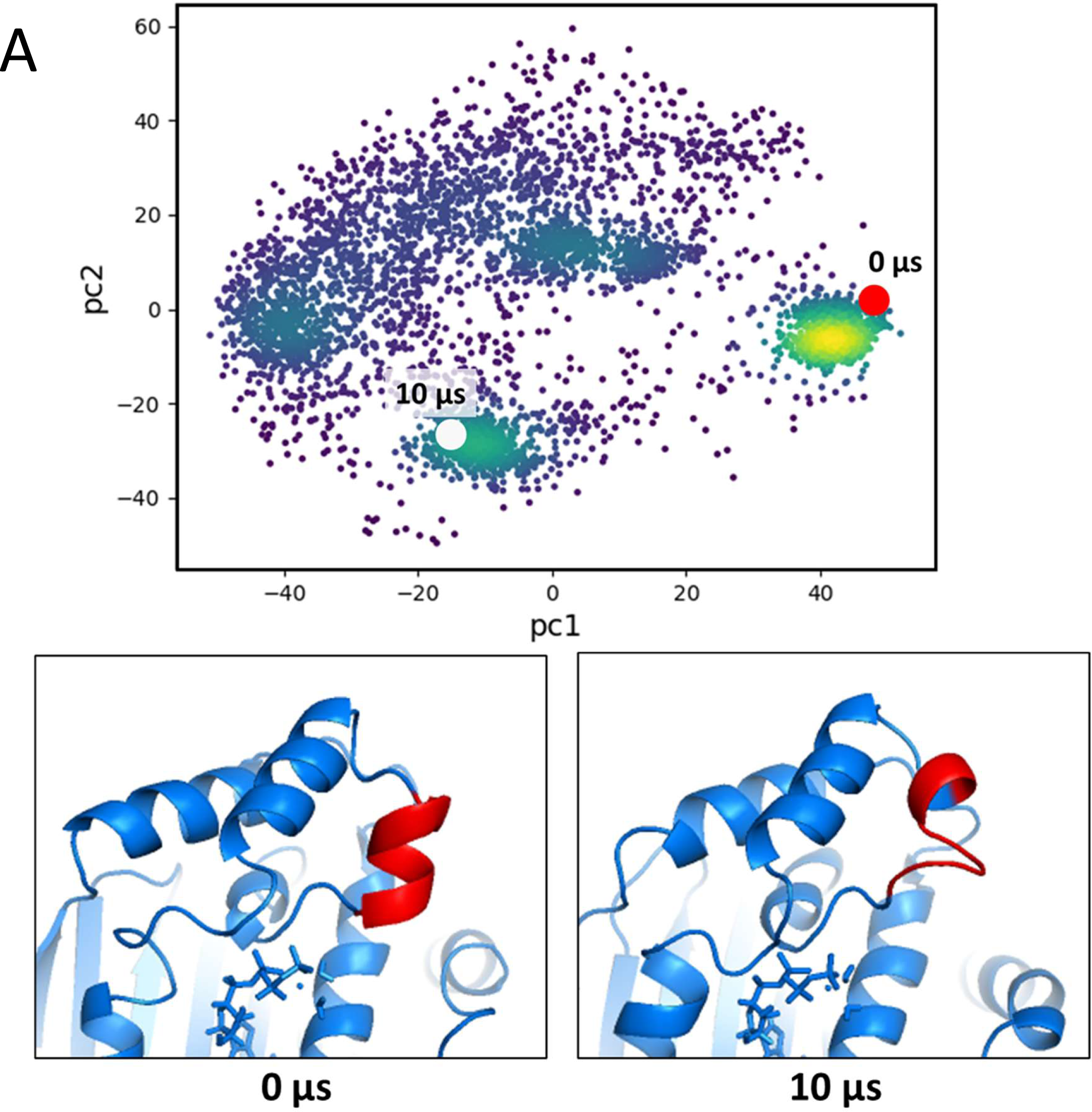
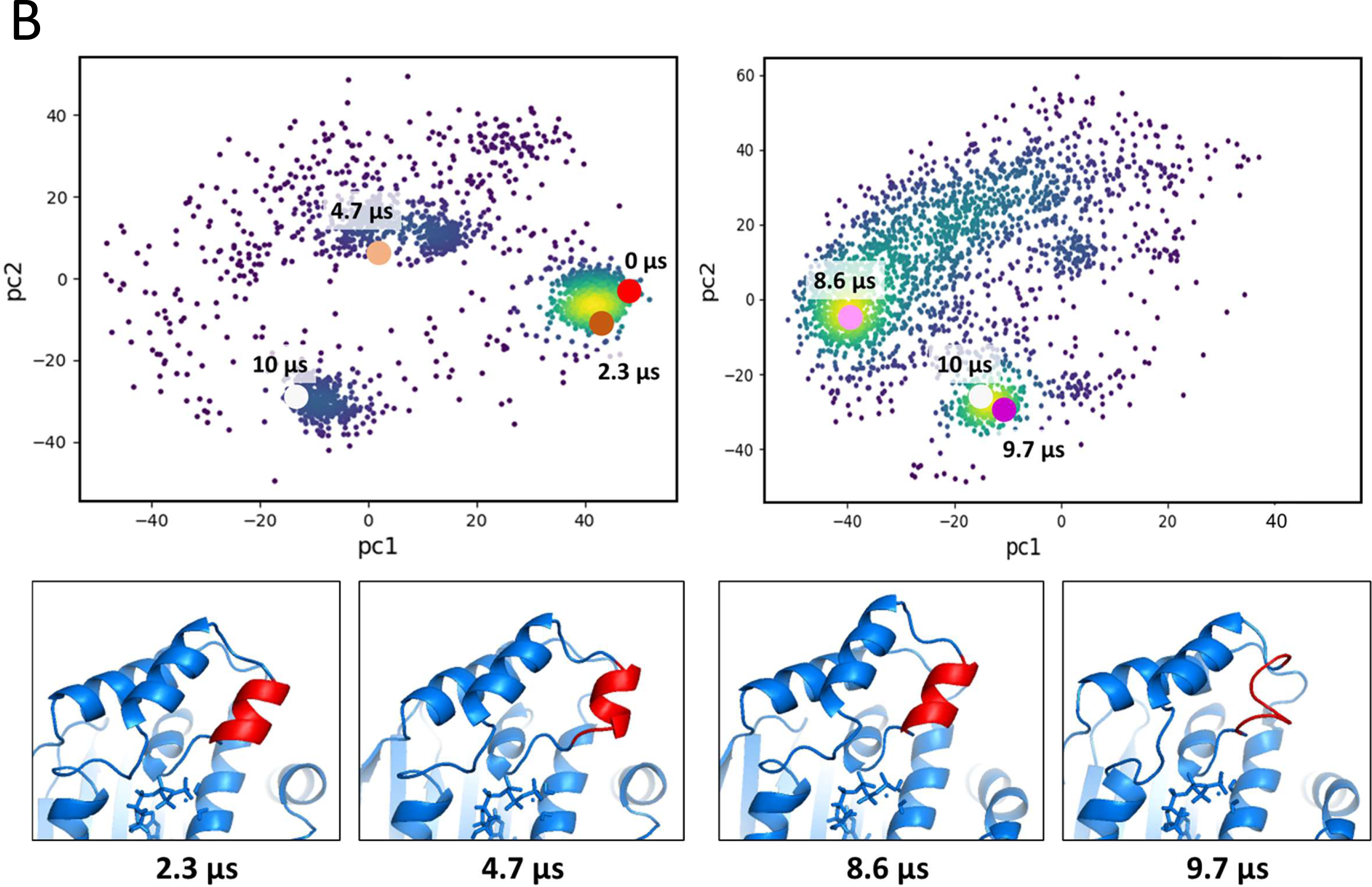
PCA of the ATP-complex structure. (A) PCA projection of the 10-μs simulation. The initial structure (0 μs) and the final structure (10 μs) of the simulation were plotted on the projection map with the red and white circles, and shown in the left and right boxes at bottom, respectively. (B) PCA map of the structures from 0 μs to 5 μs in the top left. The structures at 2.3 μs and 4.7 μs were plotted on the projection map with the brown and orange circles and shown in the two left-boxes at bottom, respectively. PCA map of structures from 6 μs to 10 μs in the top right. The structures at 8.6 μs and 9.7 μs were plotted on the projection map with the magenta and purple circles and shown in the two right-boxes at bottom, respectively. In the structure boxes, H7 segment was colored in red.

Next, the stability of the lid segment in the up-conformation was estimated from the interaction energy between the lid segment and the region facing it. Energy was estimated by binding energy analysis using the MMGBSA method (Onufriev et al., 2000; Onufriev et al., 2004). The production trajectories of the lid interaction energy exhibited larger fluctuations in the ATP-complex structure than that in the apo structure **(Figure S3)**. The 10-μs average energy of lid interaction in the ATP-complex structure was statistically bigger than that in the apo structure (-100.9±11.20 kcal/mol and -103.8±6.50 kcal/mol, p<0.001).

These results suggest that the lid segment in the ATP-complex structure fluctuated more and interacted more weakly with the facing region than that in the apo structure.

### 3.2. Effect of A107N mutation on the lid segment

To clarify effects of A107N and T101I mutations on the structure and function of the lid segment in ATP-complex, a long 10-μs simulation for the mutant structures complexed with ATP was conducted and compared with that of wild-type ATP-complex structure.

#### 3.2.1. Preparation of A107N mutant structure complexed with ATP

The X-ray structure of the A107N NTD complexed with ATP is not available. The A107N ATP-complex structure was thus modeled from the X-ray structure of the wild-type ADP-complex, similar to the modeling of the wild-type ATP-complex structure, with an *in silico* mutation of Ala107 on the H6 segment to Asn. However, as seen in the equilibration simulation of the wild-type apo and ATP-complex structures, the X-ray structures of the NTD are likely to have a crystal-packing effect on the lid structure. Therefore, 5 μs simulation was implemented for both A107N apo and ATP-complex structures, respectively, to assess the crystal-packing effect.

The 5 μs trajectory of the A107N ATP-complex showed a similar degree of conformational fluctuation on the lid segment to the equilibration trajectory of the wild-type ATP-complex structure, and both the whole structures, including lid segment, were well overlapped **(Figure S4)**. In contrast, the 5 μs trajectory of the A107N apo structure showed a large conformational change in the H7 segment (Figure 6). The H7 segment was largely displaced and was greatly unwound to form a loop conformation at around 3.5 μs. The H6 segment was similar to that of the wild-type apo structure.

**Figure 6.**
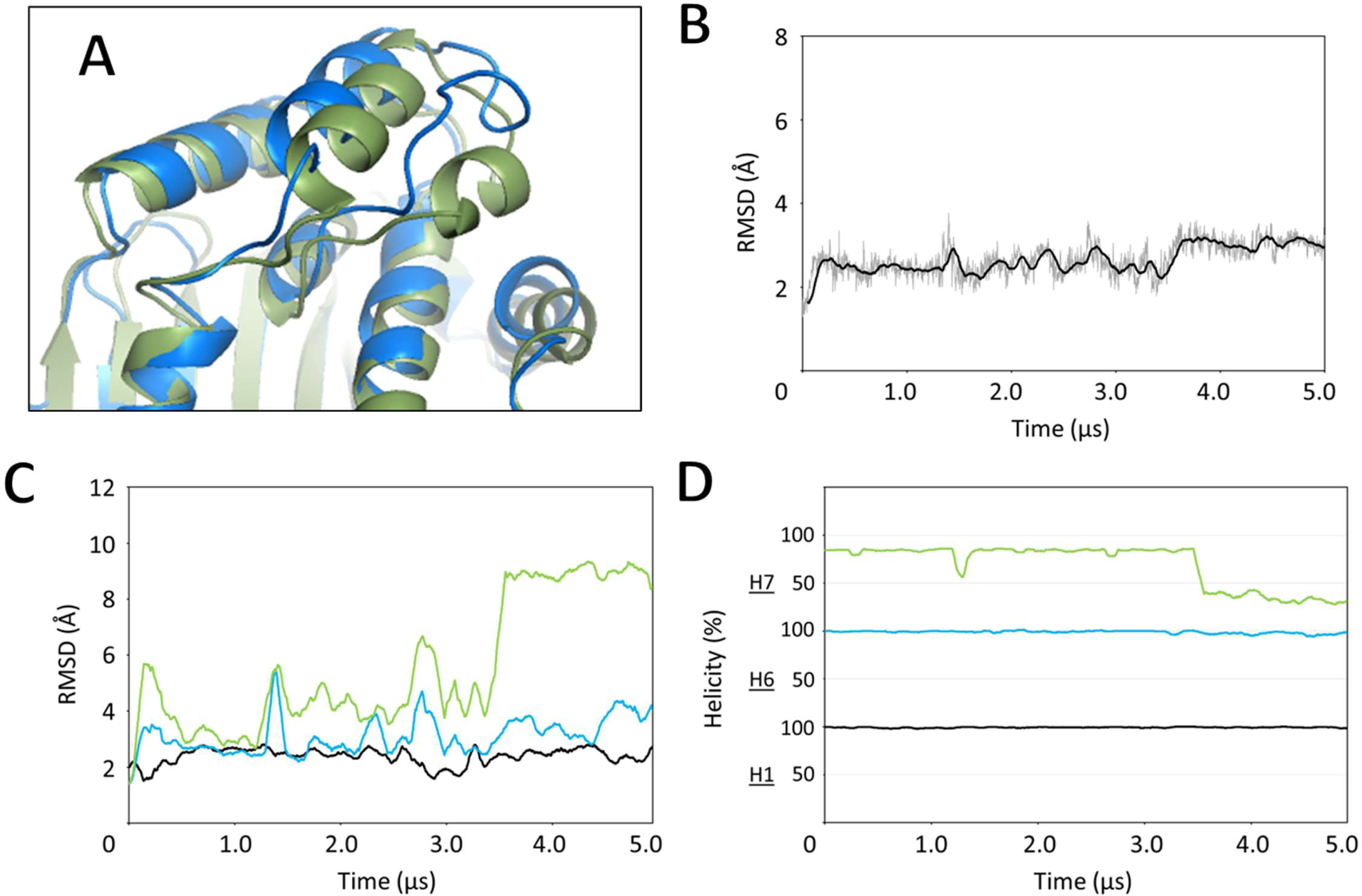
5-μs simulation for the A107N apo structure. (A) A107N apo structure at 5 μs simulation (blue) compared with the equilibrated wild-type apo structure (green), the RMSD trajectories of the entire structure (B) and the H1/H6/H7 segments (C), and (D) the helicity trajectories of the H1/H6/H7 segments. Plots of the H1/H6/H7 segments are colored in black, light blue and light green, respectively.

The large conformational difference in the H7 segment between the A107N apo and wild-type apo structures suggests that the H7 conformational change is not only due to the effect of crystal packing but also the effect of the A107N mutation in the lid segment. Therefore, the H7 segment of the A107N apo structure in a stable isolated state is most likely to adopt a conformation different from that of the wild-type apo structure. This indicated that the X-ray structure of the wild-type ADP-complex is inappropriate for use in this study to model the A107N ATP-complex structure. Thus, A107N apo structure at 5 μs in the simulation was treated as a stable A107N apo structure, which was used for an initial A107N ATP-complex structure with the addition of an ATP molecule, a Mg^2+^ ion, and water molecules for an equilibration phase of MD simulation (5 μs). During equilibration, the H7 segment was additionally displaced, and H7 unwinding continued; however, the other part was stable **(Figure S5)**. This equilibrated A107N ATP-complex structure was used for a subsequent production phase (10 μs MD).

#### 3.2.2. Structural comparison of A107N ATP-complex with wild-type ATP-complex

During production, the H7 segment was displaced, and the large decrease in H7 helicity was unchanged, while H6 helicity was retained **(Figure S6)**. The transition of the lid segment from the up- to the down-conformation was not observed.

A structural comparison of the A107N ATP-complex with the wild-type ATP-complex revealed that the H7 segment in A107N had largely departed from the region facing the lid segment **(Figure 7 A)**. The average distances between Ile117 on the H7 segment and Ile29 on the H2 segment in the facing region were 17.49±2.247 Å and 10.32±1.744 Å in the A107N ATP-complex and wild-type ATP-complex structures, respectively **(Figure 7 B,C)**. In contrast, the distance between the H6 segment and facing region, of which locations were represented by Thr101 on the H6 segment and Ile12 on the H1 segment, was not much different (A107N: 9.13±0.711 Å and wild-type: 7.07±1.073 Å) **(Figure S6)**.

**Figure 7.**
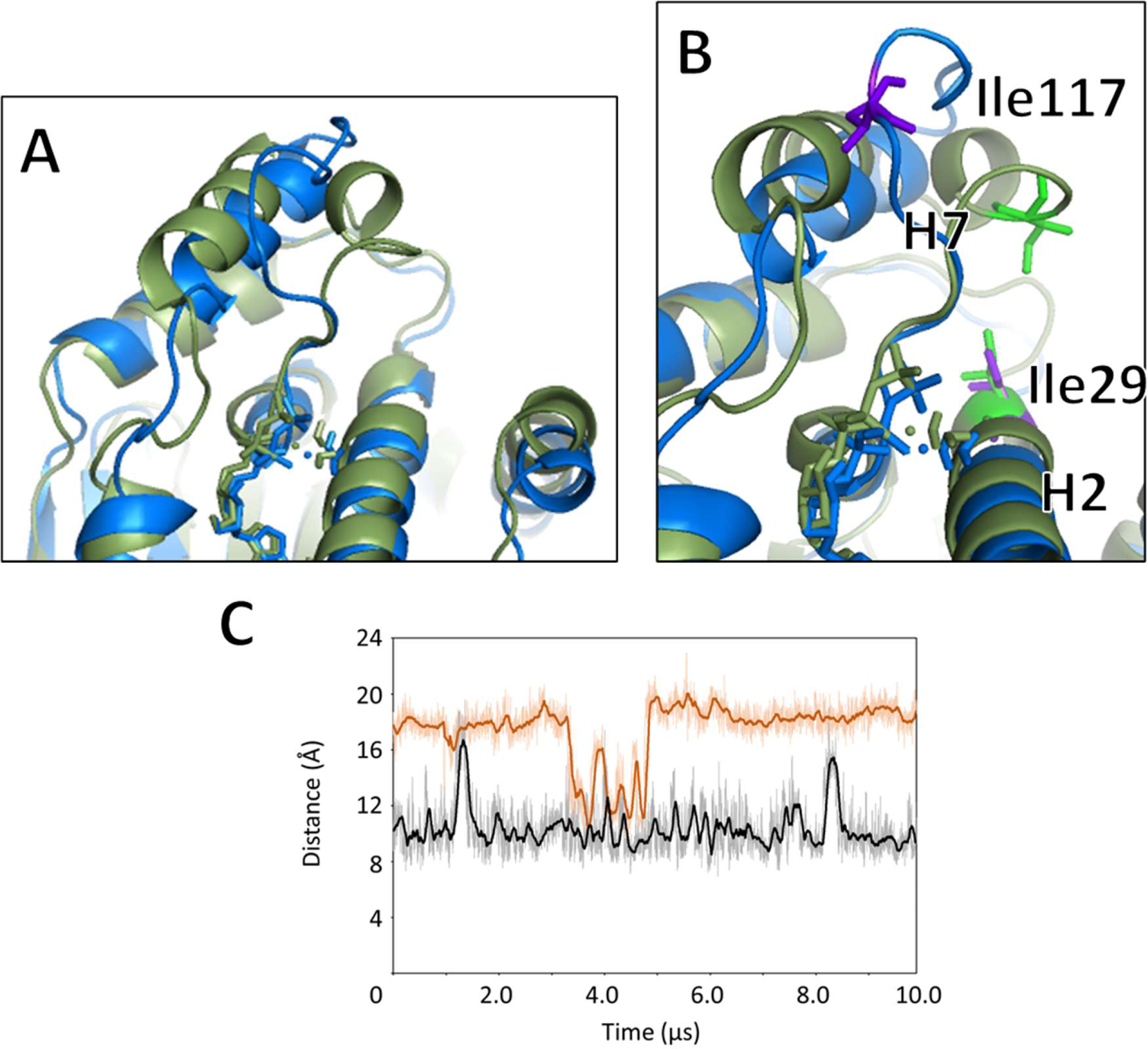
A107N ATP-complex structure after production. (A) A107N ATP-complex structure after production (blue) compared with wild-type ATP-complex structure after production (green), (B) a close-up view of the H7 segment and facing region of the A107N ATP-complex (blue) and wild-type ATP-complex structures (green). Ile117 on the H7 segment and Ile29 on the H2 segment in the A107N ATP-complex structure were colored in purple, and those in the wild-type ATP-complex structure were colored in light green, and (C) the distance trajectories between Ile29 and Ile117 (brown) and between Ile12 and Thr101 (black) in the ATP-complex structure.

This H7 leaving in the A107N ATP-complex structure should reflect the interaction energy. In fact, the average interaction energy of the lid segment in the A107N ATP-complex structure was statistically weaker than that in the wild-type ATP-complex structure (A107N: -94.6±10.46 kcal/mol and wild-type: -100.9±11.20 kcal/mol, p<0.001).

#### 3.2.3. In silico mutations in H7 segment, I117N and F120N

To determine the factor that causes unwinding of the H7 segment in the A107N ATP-complex structure, the position of the 107th residue (Ala107) in the equilibrated wild-type apo structure was visually inspected. As shown in **Figure 8 A**, the hydrophobic Ile117 is located on the H7 segment facing Ala107. It was suspected that the disturbance of this hydrophobic circumstance, due to a mutation in A107N, caused the H7 unwinding in the A107N mutant structure.

**Figure 8.**
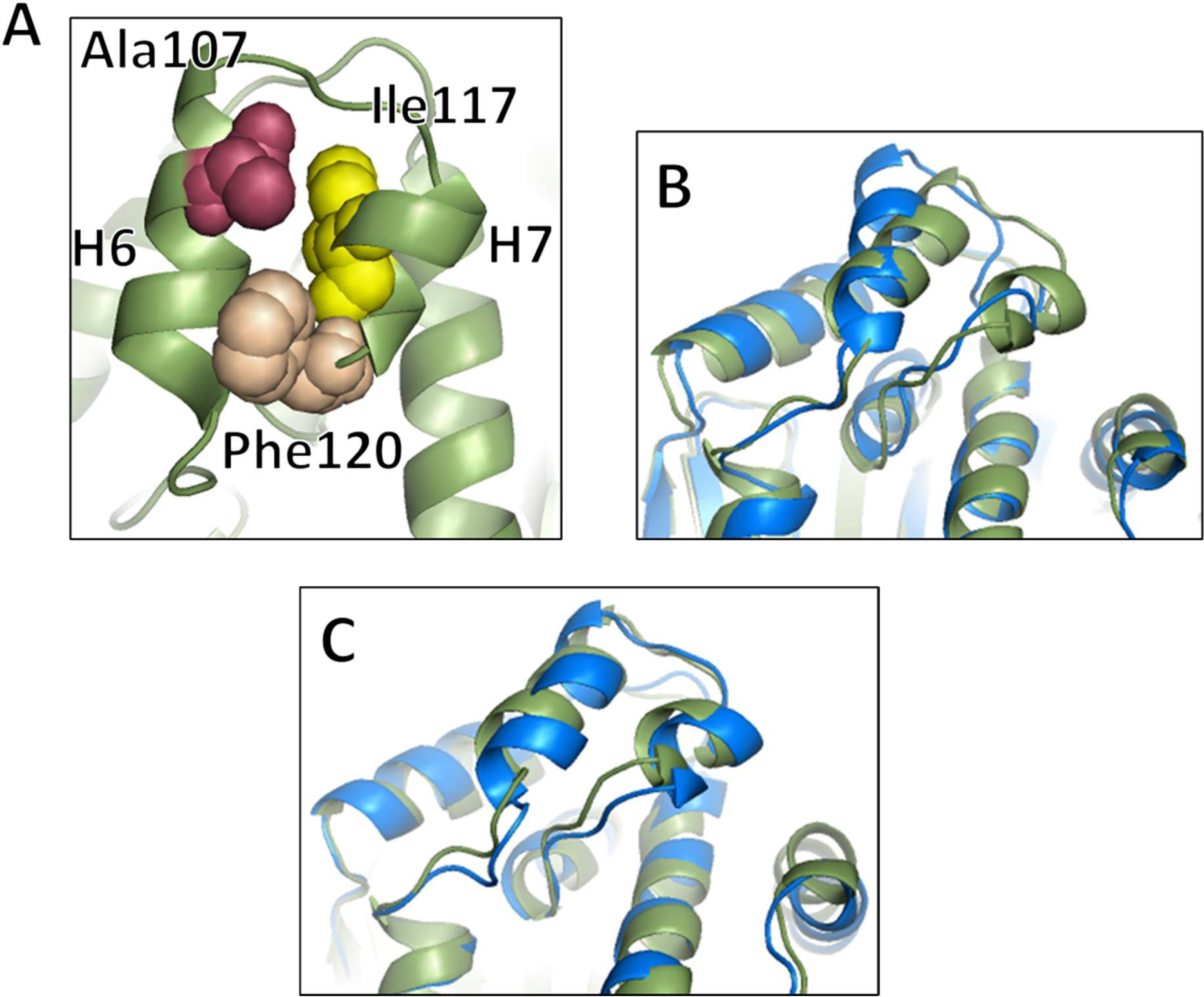
*In silico* mutation study on Ala107 neighboring residues. (A) a close-up view of Ala107 neighboring residues in the equilibrated wild-type apo structure. Ala107 on the H6 segment, Ile117 and Phe120 on the H7 segment were colored in magenta, yellow, and beige, respectively, (B) I117N apo structure after 5 μs simulation (blue) compared with wild-type apo structure after 5 μs in the production simulation (green), and (C) F120N apo structure after 5 μs simulation (blue) compared with the wild-type apo structure after 5 μs in the production simulation (green).

To examine this hypothesis, Ile117 on the H7 segment was mutated *in silico* to Asn in the equilibrated wild-type apo structure in which the H7 segment kept helicity, and subjected to 5 μs simulation. As a result, the H7 segment of the I117N apo structure after 5 μs in the trajectory was unwound and became a loop conformation **(Figure 8 B)**. Furthermore, for comparison, the hydrophobic Phe120 on the H7 segment, located slightly farther from Ala107, was mutated to Asn in the equilibrated wild-type apo structure. A 5 μs simulation for the F120N apo structure showed an unchanged structure relative to the wild-type apo structure after 5 μs in the production simulation **(Figure 8 C)**. These *in silico* mutation studies indicates that disturbance of the hydrophobic circumstance at the 107th residue is a plausible factor for H7 unwinding in the A107N mutant structure.

### 3.3. Effect of T101I mutation on the lid segment

#### 3.3.1. Preparation of T101I mutant structure complexed with ATP

As was the case in the A107N mutant structure, the T101I ATP-complex structure was modeled from the T101I apo structure after 5 μs simulation with the addition of an ATP molecule, a Mg^2+^ ion, and water molecules. This was because the X-ray structure of T101I complexed with ATP was not available; additionally, the 5 μs simulation of the T101I apo structure showed a conformational displacement and unwinding of H7 segment due to crystal-packing and mutation effects **(Figure S7)**. The T101I apo structure was mutated *in silico* using the X-ray wild-type apo structure.

This modeled T101I ATP-complex structure from the T101I apo structure was subjected to 5 μs simulation for an equilibration phase. The T101I ATP-complex structure did not change significantly during equilibration and was similar to the equilibrated wild-type ATP-complex structure, except for the H7 segment **(Figure S8)**. This equilibrated T101I ATP-complex structure was used for a production phase (10 μs simulation).

#### 3.3.2. Structural comparison of T101I ATP-complex with wild-type ATP-complex

During production, the T101I ATP-complex structure showed the displacement of the lid segment compared to that of the equilibrated T101I ATP-complex **(Figure 9 A)**. The transition of the lid segment from the up- to the down-conformation was not observed. No significant unwinding was observed for the H7 segment in the T101I ATP-complex structure after production, contrary to the case of the A107N ATP-complex. Compared to the wild-type ATP-complex structure, the position of the H6 segment was slightly displaced in the T101I ATP-complex structure, and the relative positions of the H7 segment and the facing region, the H2 segment, did not change significantly **(Figure S9)**. The average distances between Ile29 in the H2 segment and Ile117 in the H7 segment were 9.94±2.069 Å in the T101I ATP-complex and 10.32±1.744 Å in the wild-type ATP-complex structures, respectively.

**Figure 9.**
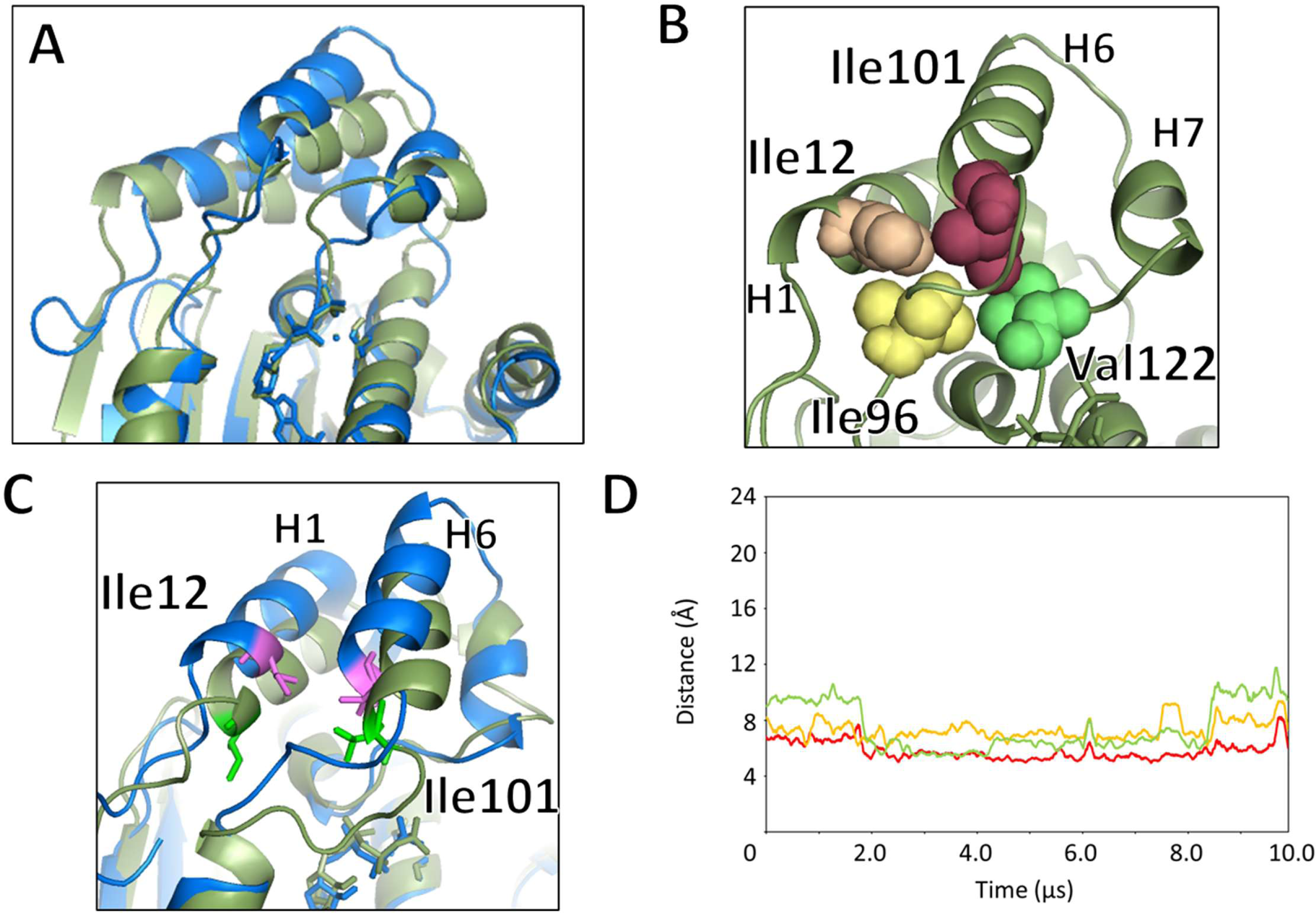
T101I ATP-complex structure after production. (A) T101I ATP-complex structure after production (blue) compared with the equilibrated T101I ATP-complex structure (green), (B) a close-up view of Ile101 neighboring residues in the T101I ATP-complex structure. Ile101 on the H6 segment, Ile12 on the H1 segment, and Ile96 and Val122 on the loop regions were colored in magenta, beige, yellow, and green, respectively, (C) a close-up view of the H6 segment and facing region of the T101I ATP-complex structure after production (blue) compared with the wild-type ATP-complex structure (green). Ile101 on the H6 segment and Ile12 on the H1 segment in the T101I ATP-complex structure were colored in purple, and those in the wild-type ATP-complex structure were colored in light green, and (D) the distance trajectories between Ile12 and Ile101 (red), between Ile96 and Ile101 (brown), and between Val122 and Ile101 (green) in the T101I ATP-complex structure during the production.

A close-up view of Ile101 on the H6 segment showed that there was a hydrophobic cluster, including Ile12 on the H1 segment and Ile96 and Val122 on the loop regions in the lid segment **(Figure 9 B)**. For example, the relative positions of Ile101 and Ile12 were closer than those of T101 and Ile12 in the wild-type ATP-complex structure **(Figure 9 C)**. The average distances between these residues were 5.81± 0.905 Å in the T101I ATP-complex and 7.07±1.073 Å in the wild-type ATP-complex structures, respectively. The relative positions of Ile101 and other hydrophobic residues (Ile96 and Val122) also retained close proximity in the T101I ATP-complex structure during production **(Figure 9 D)**. This suggests that Ile101 on the H6 segment of the T101I ATP-complex structure interacted more tightly with the facing region than with the wild-type ATP-complex structure.

These hydrophobic interactions may have stabilized the lid segment of the T101I ATP-complex structure. In fact, the average interaction energy of the lid segment in the T101I ATP-complex structure was statistically smaller than that in the wild-type ATP-complex structure (-105.3±12.43 kcal/mol and -100.9±11.20 kcal/mol, p<0.001).

## 4. Discussion

In the catalytic cycle of Hsp90, ATP hydrolysis is a key event that drives structural changes, including the interchange of the dimeric Hsp90 structure between open and closed forms. For ATP hydrolysis, the ATP-lid closure occurring in the NTD is an indispensable co-ordinated structural change. Lid closure is a phenomenon in which the upward conformation of the lid segment in the apo structure changes to a downward conformation in the ATP-complex structure along with ATP binding at the ATP-binding site. However, information regarding the atomistic mechanism of lid closure is limited. In this study, a computational biochemistry approach was applied to obtain new insights into lid closure.

A total of 15 μs MD simulations, including the equilibration and production phases, were conducted for the apo and ATP-complex structures starting from up-lid conformation, but no lid closures were observed. While previous studies using spectroscopic techniques have reported that the up-to-down transition of the lid segment occurs on the order of microseconds to milliseconds (Henot et al., 2022; Schubert et al., 2021; Schulze et al., 2016), the early microsecond timescale conducted in this study has been insufficient to monitor this process. However, a very early event, i.e., *a sign*, of lid closure has been captured in this study.

In the simulation of the ATP-complex structure, the lid segment fluctuated more than in the apo structure, and unwinding of the H7 segment was observed in the lid segment of the ATP-complex structure. This conformational instability of the lid segment energetically weakened the interaction with its facing region and caused the lid to leave the facing region, suggesting that the up-to-down transition was triggered by the conformational instability of the lid segment, particularly the H7 segment. In comparing the ATP-complex structure with the apo structure, this instability may be due to a sterically pushed-up loop in the lid segment by neighboring ATP γ-phosphate.

Furthermore, the simulation of the A107N ATP-complex structure showed a much clearer H7 unwinding and leaving due to the loss of helical packing with the H6 segment.

In the simulation of the ATP-complex structures of A107N and T101I mutants that had mutations in the lid segment and altered ATP hydrolytic activity, the interaction energy between the lid segment and facing regions was ranked in the order A107N (-94.6±10.46 kcal/mol) > wild-type (-100.9±11.20 kcal/mol) > T101I (-105.3±12.43 kcal/mol), which coincided well with the order of ATP hydrolytic activity (i.e., A107N > wild-type > T101I). In addition, ATP-spiking experiments in a fluorescence resonance energy transfer (FRET) study showed that conformational changes of the lid segment in the apo structure spiked by ATP occurred promptly in the order of A107N > wild-type > T101I (Schulze et al., 2016). This was in good agreement with the simulation results.

The kinetic mechanism of lid closure via ATP binding has not yet been clarified regarding whether conformational selection or induced fit is dominant. A recent NMR study in human Hsp90ɑ reports that the conformational interchange of the lid segment between up- and down-conformations occurs in the apo structure, and the populations of up- and down-conformations are estimated to be 96.8±0.1 % and 3.2±0.1 %, respectively (Henot et al., 2022). It is therefore proposed that lid closure in human Hsp90ɑ is dominated by the conformational selection mechanism. In this study, observation of the sign of the lid leaving the ATP-binding site, in simulations of the wild-type and A107N ATP-complex structures through H7 unwinding, supports the conformation selection mechanism because the lid leaving after ATP binding should decrease the population of the up-conformation and increase the intermediate- or down-conformation in the open form of Hsp90. FRET studies have reported that an intermediate-conformation of the lid segment exists between the up- and down-conformations (Schubert et al., 2021; Schulze et al., 2016). In addition, the direct interaction with ATP γ-phosphate was not observed in the simulation of the ATP-complex structure, except for Lys98. Lys98 continuously interacted with ATP γ-phosphate, but this interaction may just hold the phosphate group in an appropriate position in the ATP-binding site supportively. No other direct interaction between the γ-phosphate and lid segment, which should be required for the mechanism of induced fit, was observed in the ATP-complex structure.

In conclusion, our simulations suggest that lid closure from the up- to down-conformation is triggered by lid leaving accompanying the conformational instability of the lid segment, including H7 unwinding. However, as previously reported, our simulations failed to reproduce the intermediate or downward conformations of the lid segment. Other critical interactions and/or conformational changes in the intermediate- or down-conformation states may regulate the up-to-down transition of the lid segment. Also, other effects on altering ATP hydrolytic activity of A107N and T101I mutants may exist in these states. Therefore, further studies are warranted to clarify lid function and to elucidate the mechanism of the catalytic cycle of Hsp90.

## Abbreviations

FRET: fluorescence resonance energy transfer
Hsp90: 90 kDa heat shock protein
MD: molecular dynamics
MMGBSA: molecular mechanics/generalized Born surface area
NTD: N-terminal domain; RMSD, root-mean squared deviation

## References

Ali, M. M., Roe, S. M., Vaughan, C. K., Meyer, P., Panaretou, B., Piper, P. W., Prodromou, C. & Pearl, L. H. (2006). Crystal structure of an Hsp90-nucleotide-p23/Sba1 closed chaperone complex. Nature, 440, 1013–1017.

Alln, O., Nilsson, L. & Villa, A. (2012). Magnesium ion-water coordination and exchange in biomolecular simulations. Journal of Chemical Theory and Computation, 8, 1493–1502.

Andrea, T. A., Swope, W. C. & Andersen, H. C. (1983). The role of long ranged forces in determining the structure and properties of liquid water. Journal of Chemical Physics, 79, 4576−4584.

Berendsen, H. J. C., Postma, J. P. M., van Gunsteren, W. F., DiNola, A. & Haak, J. R. (1984). Molecular dynamics with coupling to an external bath. Journal of Chemical Physics, 81, 3684−3690.

Biebl, M. M., & Buchner, J. (2019). Structure, function, and regulation of the Hsp90 machinery. Cold Spring Harbor Perspectives in Biology, 11/9/a034017.

Bryce, R. (2008). AMBER parameter database. University of Manchester, Manchester, UK. http://amber.manchester.ac.uk/index.html (accessed on 1 March 2024)

Case, D. A., Aktulga, H. M., Belfon, K., Ben-Shalom, I. Y., Berryman, J. T., Brozell, S. R., Cerutti, D. S., Cheatham, III, T. E., Cisneros, G. A., Cruzeiro, V. W. D., Darden, T. A., Duke, R. E., Giambasu, G., Gilson, M. K., Gohlke, H., Goetz, A. W., Harris, R., Izadi, S., Izmailov, S. A., Kasavajhala, K., Kaymak, M. C., King, E., Ko-valenko, A., Kurtzman, T., Lee, T. S., LeGrand, S., Li, P., Lin, C., Liu, J., Luchko, T., Luo, R., Machado, M., Man, V., Manathunga, M., Merz, K. M., Miao, Y., Mikhailovskii, O., Monard, G., Nguyen, H., O’Hearn, K. A., Onufriev, A., Pan, F., Pantano, S., Qi, R., Rahnamoun, A., Roe, D. R., Roitberg, A., Sagui, C., Schott-Verdugo, S., Shajan, A., Shen, J., Simmerling, C. L., Skrynnikov, N. R., Smith, J., Swails, J., Walker, R. C., Wang, J., Wang, J., Wei, H., Wolf, R. M., Wu, X., Xiong, Y., Xue, Y., York, D. M., Zhao, S. & Kollman, P. A. (2022). *Amber* 2022, University of California, San Francisco, USA.

Darden, T., York, D. & Pedersen, L. (1993). Particle mesh Ewald: An N·log(N) method for Ewald sums in large systems. Journal of Chemical Physics, 98, 10089−10092.

Dolinsky, T. J., Czodrowski, P., Li, H., Nielsen, J. E., Jensen, J. H., Klebe, G. & Baker, N. A. (2007). PDB2PQR: expanding and upgrading automated preparation of biomolecular structures for molecular simulations. Nucleic Acids Research, 35, W522−W525.

Essmann, U., Perera, L., Berkowitz, M. L., Darden, T., Lee, H. & Pedersen, L. G. (1995). A smooth particle mesh Ewald method. Journal of Chemical Physics, 103, 8577−8593.

Goetz, A. W., Williamson, M. J., Xu, D., Poole, D., Le Grand, S., Walker, R. C. (2012). Routine microsecond molecular dynamics simulations with AMBER - Part I: Generalized Born. Journal of Chemical Theory and Computation, 8, 1542–1555.

Gohda, K. (2022). Conformational analysis of the loop-to-helix transition of the α-helix3 plastic region in the N-terminal domain of human Hsp90α by a computational biochemistry approach. Journal of Chemical Information and Modeling, 62, 5699– 5714.

Gohlke, H. & Case, D. A. (2004). Converging free energy estimates, MM-PB(GB)SA studies on the protein-protein complex Ras-Raf. Journal of Computational Chemistry, 25, 238–250.

Gohlke, H., Kiel, C. & Case, D. A. (2003). Insights into protein-protein binding by binding free energy calculation and free energy decomposition for the Ras-Raf and Ras-RalGDS complexes. Journal of Molecular Biology, 330, 891–913.

Hawkins, G. D., Cramer, C. J. & Truhlar, D. G. (1995) .Pairwise solute descreening of solute charges from a dielectric medium. Chemical Physics Letter, 246, 122–129.

Hawkins, G. D., Cramer, C. J. & Truhlar, D. G. (1996). Parametrized models of aqueous free energies of solvation based on pairwise descreening of solute atomic charges from a dielectric medium. Journal of Physical Chemistry, 100, 19824–19839.

Henot, F., Rioual, E., Favier, A., Macek, P., Crublet, E., Josso, P., Brutscher, B., Frech, M., Gans, P., Loison, C. & Boisbouvier, J. (2022). Visualizing the transiently populated closed-state of human HSP90 ATP binding domain. Nature Communications, 13, 7601.

Hoy, S. M. (2022). Pimitespib: first approval. Drugs, 82, 1413–1418.

Ibragimova, G. T. & Wade, R. C. (1998). Importance of explicit salt ions for protein stability in molecular dynamics simulation. Biophysical Journal, 74, 2906−2911.

Jorgensen, W. L., Chandrasekhar, J., Madura, J. & Klein, M. L. (1983). Comparison of simple potential functions for simulating liquid water. Journal of Chemical Physics, 79, 926−935.

Kabsch, W. & Sander, C. (1983). Dictionary of protein secondary structure: Pattern recognition of hydrogen-bonded and geometrical features. Biopolymers, 22, 2577−2637.

Le Grand, S., Goetz, A. W. & Walker, R. C. (2013). SPFP: Speed without compromise - a mixed precision model for GPU accelerated molecular dynamics simulations. Computer Physics Communications, 184, 374–380.

Li, J. & Buchner, J. (2013). Structure, function and regulation of the hsp90 machinery. Biomedical Journal, 36, 106–117.

Li, J., Sun, L., Xu, C., Yu, F., Zhou, H., Zhao, Y., Zhang, J., Cai, J., Mao, C., Tang, L., Xu, Y. & He, J. (2012). Structure insights into mechanisms of ATP hydrolysis and the activation of human heat-shock protein 90. Acta Biochim et Biophys Sinica, 44, 300–306.

Mader, S. L., Lopez, A., Lawatscheck, J., Luo, Q., Rutz, D. A., Gamiz-Hernandez, A. P., Sattler, M., Buchner, J. & Kaila, V. R. I. (2020). Conformational dynamics modulate the catalytic activity of the molecular chaperone Hsp90. Nature Communications, 11, 1410.

Maier, J. A., Martinez, C., Kasavajhala, K., Wickstrom, L., Hauser, K. E. & Simmerling, C. (2015). ff14SB, Improving the accuracy of protein side chain and backbone parameters from ff99SB. Journal of Chemical Theory and Computation, 11, 3696−3713.

Meagher, K. L., Redman, L. T. & Carlson, H. A. (2003). Development of polyphosphate parameters for use with the AMBER force field. Journal of Computational Chemistry, 24, 1016–1025.

Meyer, P., Prodromou, C., Hu, B., Vaughan, C., Roe, S. M., Panaretou, B., Piper, P. W. & Pearl, L. H. (2003). Structural and functional analysis of the middle segment of hsp90: implications for ATP hydrolysis and client protein and cochaperone interactions. Molecular Cell, 11, 647−658.

Miller, B. R., Dwight, T. M., Swails, J. M., Homeyer, N., Gohlke, H. & Roitberg, A. E. (2012). MMPBSA.py: An efficient program for end-state free energy calculations. Journal of Chemical Theory and Computation, 8, 3314–3321.

Nathan, D. F. & Lindquist, S. (1995). Mutational analysis of Hsp90 function: interactions with a steroid receptor and a protein kinase. Molecular Cell Biology, 15, 3917–25.

Nemoto, T., Ohara-Nemoto, Y., Ota, M., Takagi, T. & Yokoyama, K. (1995). Mechanism of dimer formation of the 90-kDa heat-shock protein. European Journal of Biochemistry, 233, 1−8.

Onufriev, A., Bashford, D. & Case, D. A. (2000). Modification of the generalized Born model suitable for macromolecules. Journal of Physical Chemistry B, 104, 3712– 3720.

Onufriev, A., Bashford, D. & Case, D. A. (2004). Exploring protein native states and large-scale conformational changes with a modified generalized Born model. Proteins, 55, 383–394.

Prodromou, C. (2016). Mechanisms of Hsp90 regulation. Biochemical Journal, 473, 2439–2452.

Prodromou, C. & Bjorklund, D. M. (2022). Advances towards understanding the mechanism of action of the Hsp90 complex. Biomolecules, 12, 600.

Prodromou C., Panaretou, B., Chohan, S., Siligardi, O’Brien, G. R., Ladbury, J. E., Roe, S. M., Piper, P. W. & Pearl, L. H. (2000). The ATPase cycle of Hsp90 drives a molecular ‘clamp’ via transient dimerization of the N-terminal domains. EMBO Journal, 19, 4383–4392.

Prodromou, C., Roe, S. M., O’Brien, R., Ladbury, J. E., Piper, P. W. & Pearl, L. H. (1997a). Identification and structural characterization of the ATP/ADP-binding site in the Hsp90 molecular chaperone. Cell, 90, 65−75.

Prodromou, C., Roe, S. M., Piper, P. W. & Pearl, L. H. (1997b). A molecular clamp in the crystal structure of the N-terminal domain of the yeast Hsp90 chaperone. Nature Structural Biology, 4, 477–482.

Reidy, M. & Masison, D. C. (2020). Mutations in the Hsp90 N domain identify a site that controls dimer opening and expand human Hsp90α function in yeast. Journal of Molecular Biology, 432, 4673–4689.

Roe, D. R. & Cheatham, T. E., III. (2013). PTRAJ and CPPTRAJ: Software for processing and analysis of molecular dynamics trajectory data. Journal of Chemical Theory and Computation, 9, 3084–3095.

Richter, K., Reinstein, J. & Buchner, J. (2002). N-terminal residues regulate the catalytic efficiency of the Hsp90 ATPase cycle. Journal of Biological Chemistry, 277, 44905–10.

Ryckaert, J.-P., Ciccotti, G. & Berendsen, H. J. C. (1977). Numerical integration of the cartesian equations of motion of a system with constraints: Molecular dynamics of n-alkanes. Journal of Computational Physics, 23, 327−341.

Salomon-Ferrer, R., Goetz, A. W., Poole, D., Le Grand, S. & Walker, R. C. (2013). Routine microsecond molecular dynamics simulations with AMBER - Part II: Particle Mesh Ewald. Journal of Chemical Theory and Computation, 9, 3878–3888.

Schubert, J., Schulze, A., Prodromou, C. & Neuweiler, H. (2021). Two-colour single-molecule photoinduced electron transfer fluorescence imaging microscopy of chaperone dynamics. Nature Communications, 12, 6964.

Schulze, A., Beliu, G., Helmerich, D. A., Schubert, J., Pearl, L. H., Prodromou, C. & Neuweiler, H. (2016). Cooperation of local motions in the Hsp90 molecular chaperone ATPase mechanism. Nature Chemical Biology, 12, 628–635.

Siligardi, G., Hu, B., Panaretou, B., Piper, P. W., Pearl, L. H. & Prodromou, C. (2004). Co-chaperone regulation of conformational switching in the Hsp90 ATPase cycle. Journal of Biological Chemistry, 279, 51989–98.

Stachowski, T. R. & Fischer, M. (2022). Large-scale ligand perturbations of the protein conformational landscape reveal state-specific interaction hotspots. Journal of Medicinal Chemistry, 65, 13692−13704.

Tsui, V. & Case, D. A. (2001). Theory and applications of the generalized Born solvation model in macromolecular simulations. Biopolymers, 56, 275–291.

Wei, H., Zhang Y., Jia Y., Chen, X., Niu, T., Chatterjee, A., He, P. & Hou, G. (2024). Heat shock protein 90: biological functions, diseases, and therapeutic targets. MedComm, 5, e470.

Xiao Y. & Liu Y. (2020). Recent advances in the discovery of novel HSP90 inhibitors: An update from 2014. Current Drug Targets, 21, 302–317.

