## Supplemental Figures S1 to S9 for "Understanding of ATP-lid conformational dynamics in the N-terminal domain of Hsp90 and its mutants by use of a computational biochemistry approach"

Keigo Gohda
Computer-aided Molecular Modeling Research Center, Kansai (CAMM-Kansai)
3-32-302, Tsuto-Otsuka, Nishinomiya 663-8241, Japan

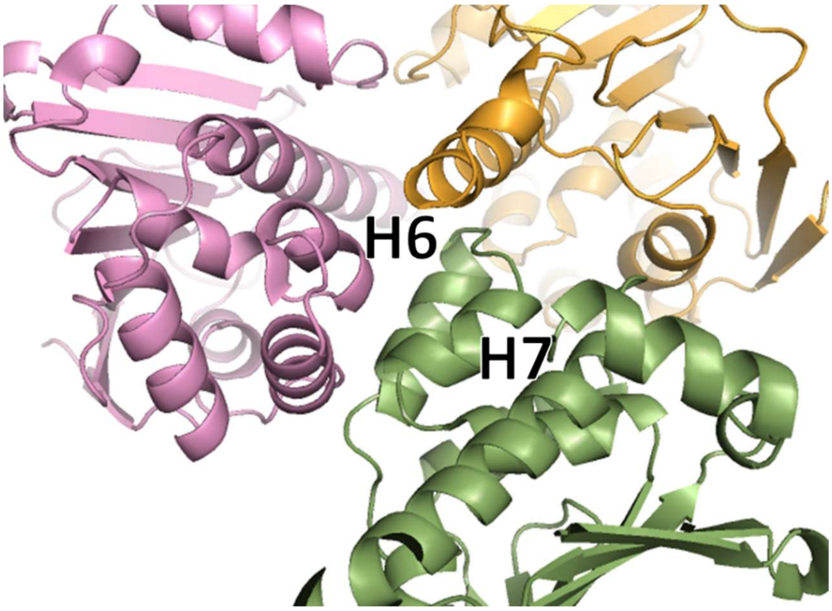


**Figure S1.** Crystal packing of lid segment in the X-ray apo structure of the NTD (green) (1AH6; Prodromou et al., 1997b). Structures in orange and magenta are symmetry-related structures.

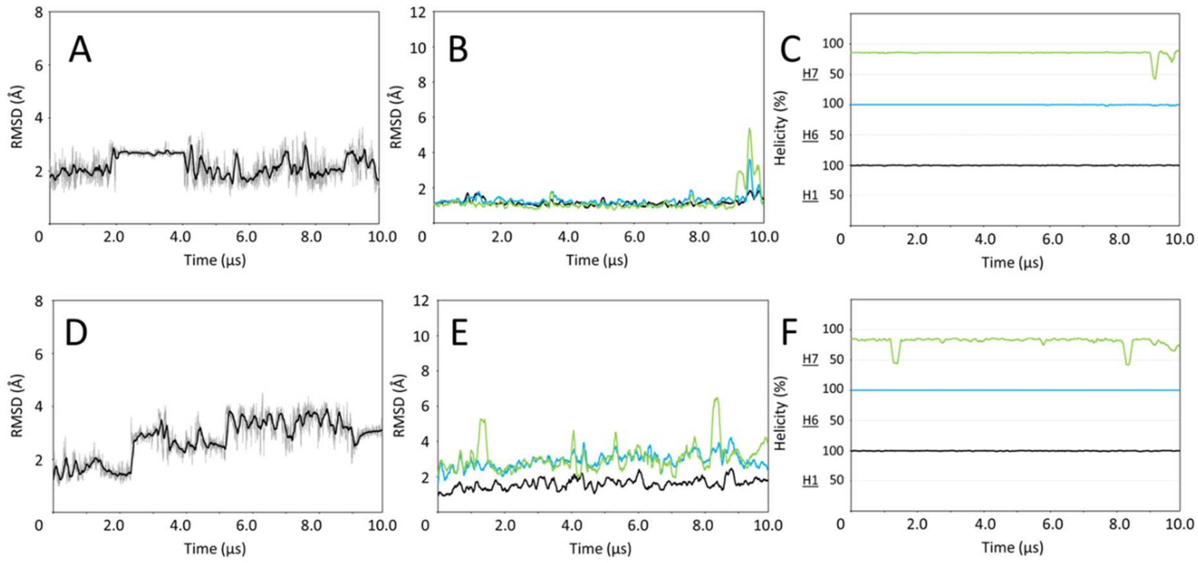


**Figure S2.** MD production for the apo and ATP-complex structures. The RMSD trajectories of the entire structure and the H1/H6/H7 segments, and the helicity trajectories of the H1/H6/H7 segments. (A-C) apo structure and (D-F) ATP-complex structure. Plots of the H1/H6/H7 segments are colored in black, light blue and light green, respectively.

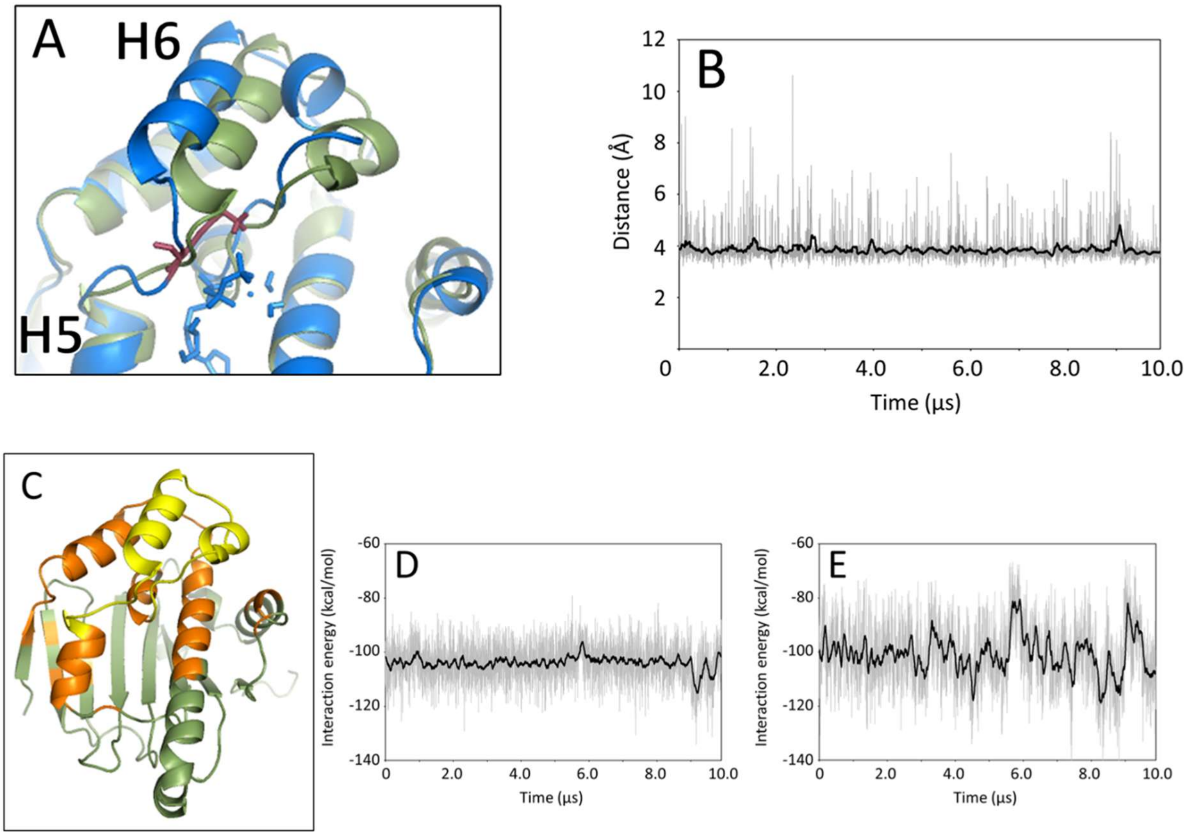


**Figure S3.** MD production for the ATP-complex structure. (A) a close-up view of the side-chain atoms of Lys98 (red) on the lid loop between the H5 and H6 segments in the ATP-complex structure after production (blue) overlapped to the apo structure after production (green), and (B) the trajectory of distance between Lys98 and ATP γ-phosphate in the ATP-complex structure, (C) "imaginary" facing regions (orange) to the lid segment (yellow) in apo structure after production. The facing region is changeable in the simulation because of the movement of the lid segment. Therefore, the facing region is denoted as imaginary facing region. The residues located within 8 Å distance from the lid segment were colored as the facing region. The trajectories of the interaction energy between the lid segment and facing regions in (D) the apo structure and (E) the ATP-complex structure, respectively.

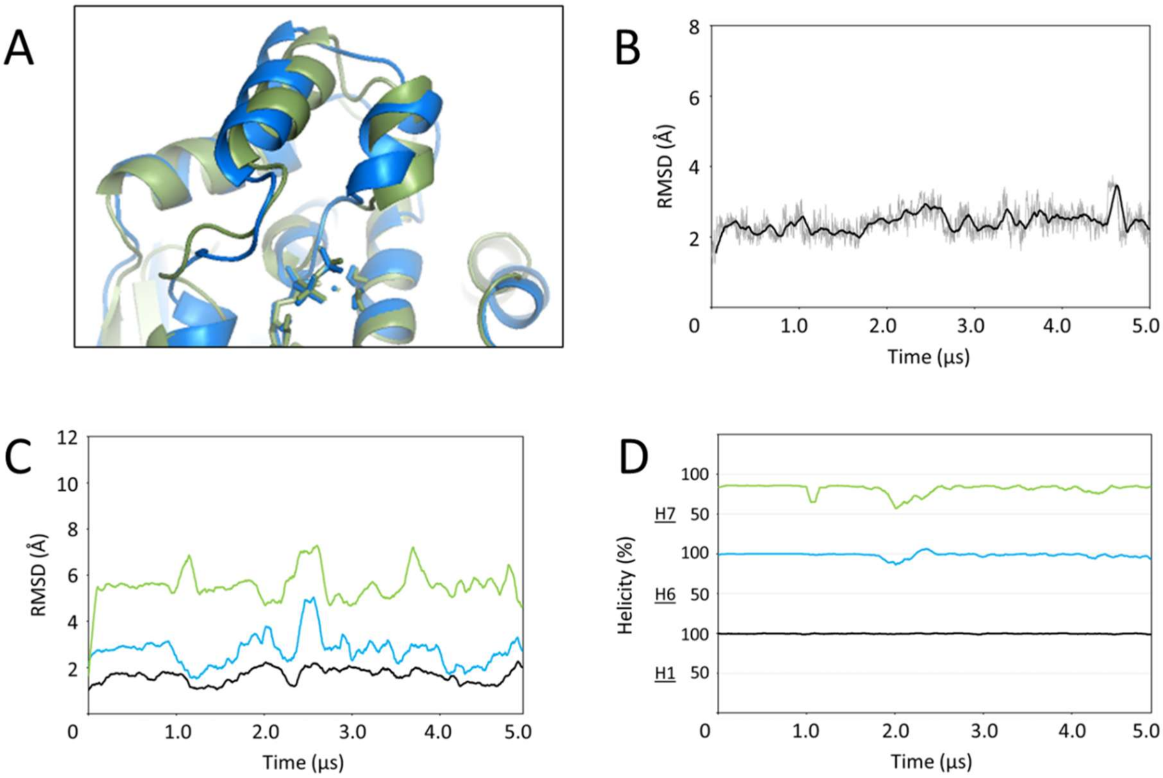


**Figure S4**. 5-μs simulation for the A107N ATP-complex structure. (A) A107N ATP-complex structure at 5 μs simulation (blue) compared with the equilibrated wild-type ATP-complex structure (green), and (B) the RMSD trajectory of the entire structure, and (C) the RMSD trajectory and (D) the helicity trajectory of the H1/H6/H7 segments. Plots of the H1/H6/H7 segments are colored in black, light blue and light green, respectively.

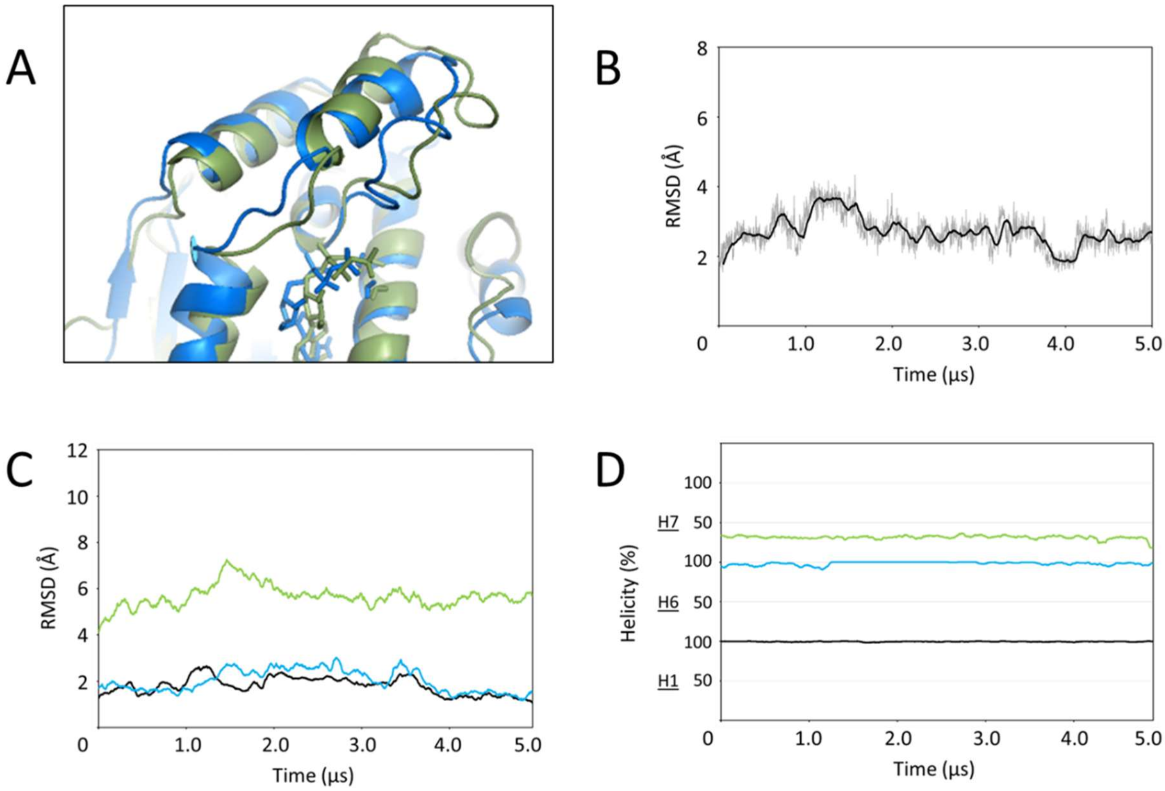


**Figure S5.** MD equilibration for the A107N ATP-complex structure. (A) A107N ATP-complex structure after equilibration (blue) compared with the initial A107N ATP-complex structure (green), and (B) the RMSD trajectory of the entire structure, and (C) the RMSD trajectory and (D) the helicity trajectory of the H1/H6/H7 segments. Plots of the H1/H6/H7 segments are colored in black, light blue and light green, respectively.

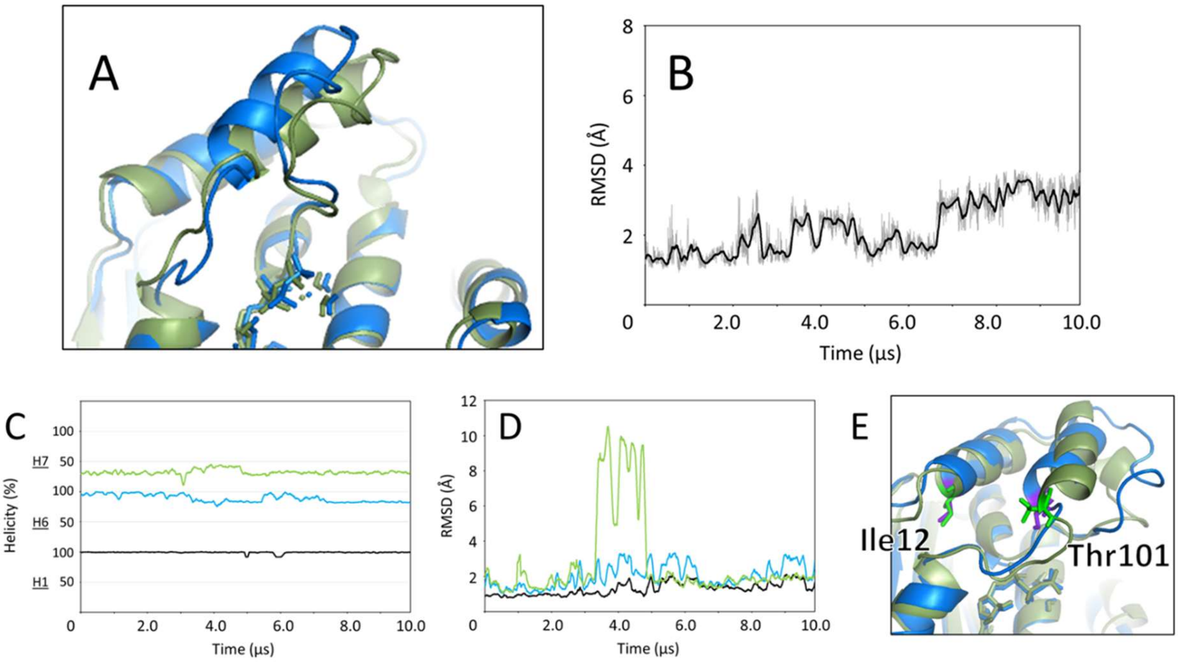


**Figure S6.** MD production for the A107N ATP-complex structure. (A) A107N ATP-complex structure after production (blue) compared with the equilibrated A107N ATP-complex structure (green), and the RMSD trajectories of (B) the entire structure and (C) the H1/H6/H7 segments, and (D) the helicity trajectories of the H1/H6/H7 segments and (E) a close-up view of the H6 segment and facing region of the A107N ATP-complex structure (blue) and the wild-type ATP-complex structure after production (green). Thr101 on the H6 segment and Ile12 on the H1 segment in the A107N ATP-complex structure were colored in purple, and those in wild-type ATP-complex structure were colored in light green. Plots of the H1/H6/H7 segments are colored in black, light blue and light green, respectively.

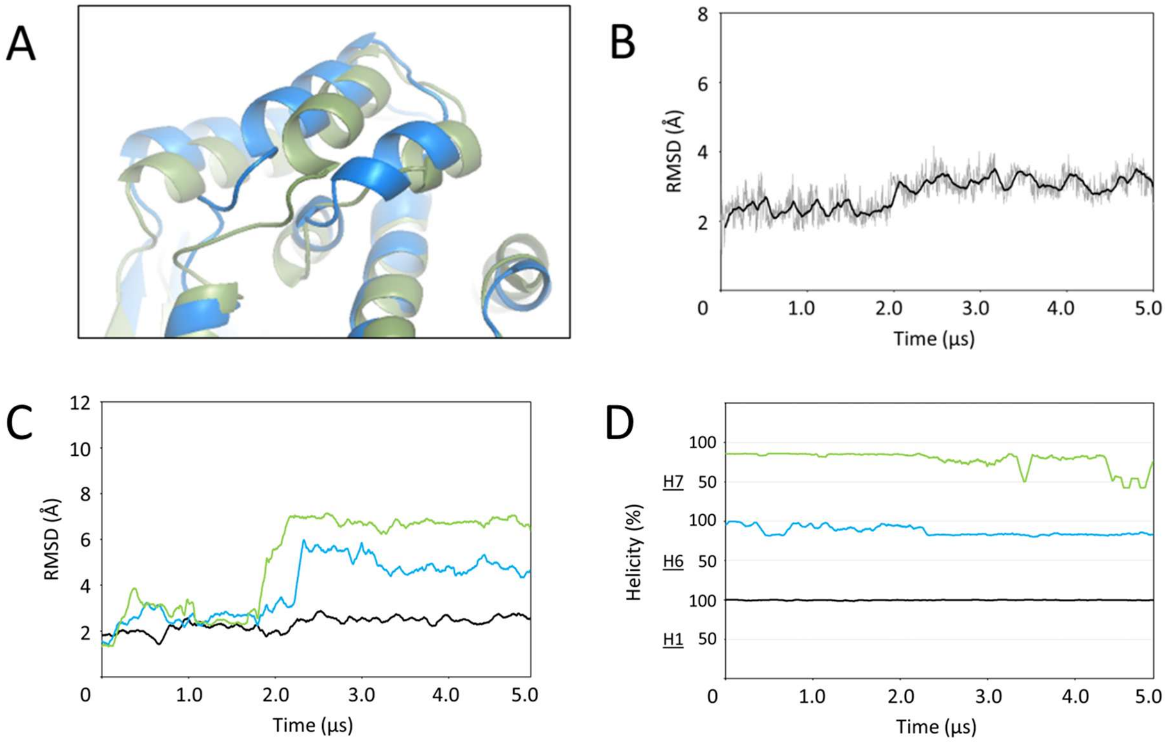


**Figure S7.** 5-μs simulation for the T101I apo structure. (A) T101I apo structure at 5 μs simulation (blue) compared with the equilibrated wild-type apo structure (green), and (B) the RMSD trajectory of the entire structure, and (C) the RMSD trajectory and (D) the helicity trajectories of the H1/H6/H7 segments. Plots of the H1/H6/H7 segments are colored in black, light blue and light green, respectively.

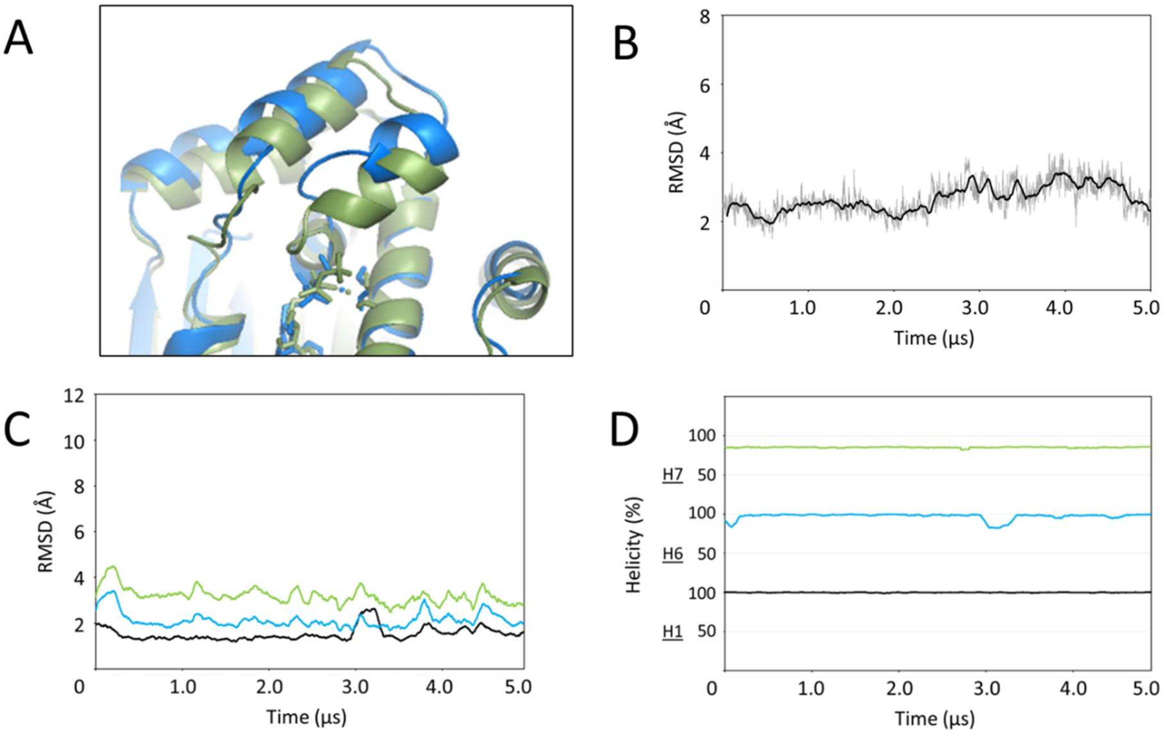


**Figure S8.** MD equilibration for the T101I ATP-complex structure. (A) T101I ATP-complex structure after equilibration (blue) compared with the initial T101I ATP-complex structure (green) and (B) the RMSD trajectory of the entire structure, and (C) the RMSD trajectory and (D) the helicity trajectories of the H1/H6/H7 segments. Plots of the H1/H6/H7 segments are colored in black, light blue and light green, respectively.

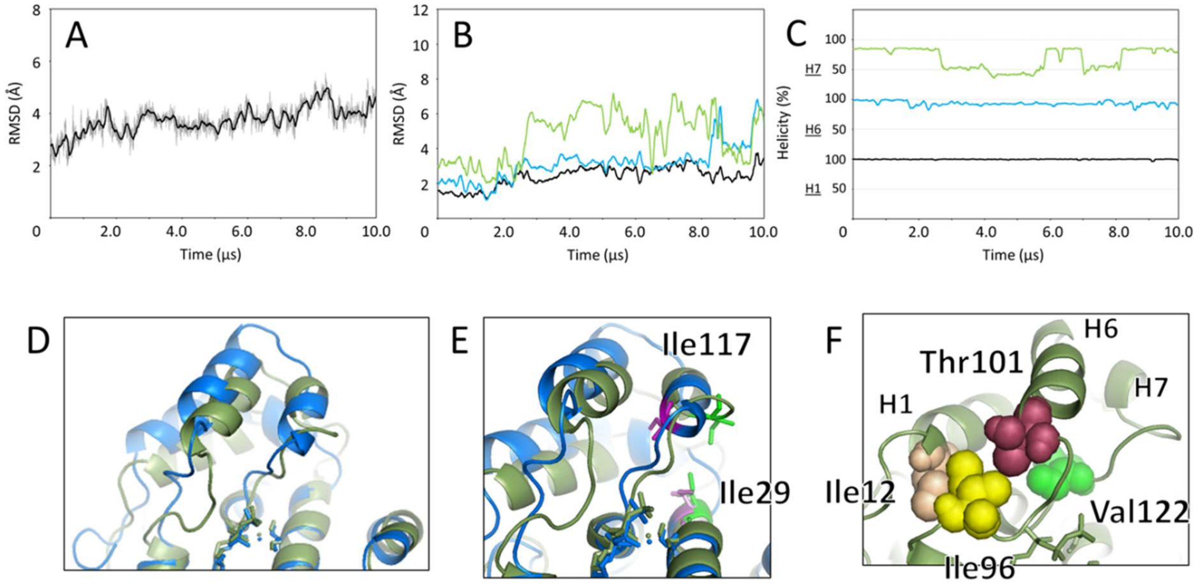


**Figure S9.** T101I ATP-complex structure after production. The RMSD trajectories of (A) the entire structure and (B) the H1/H6/H7 segments, and (C) the helicity trajectories of the H1/H6/H7 segments of the T101I ATP-complex structure during production (Plots of the H1/H6/H7 segments are colored in black, light blue and light green, respectively). (D) T101I ATP-complex structure after production (blue) compared with the wild-type ATP-complex structure (green). (E) a close-up view of the H7 segment and facing region. Ile117 on the H7 segment and Ile29 on the H2 segment in the T101I ATP-complex structure were colored in purple, and those in the wild-type ATP-complex structure were colored in light green. (F) a close-up view of Thr101 neighboring residues in the wild-type ATP-complex structure. Thr101 on the H6 segment, Ile12 on the H1 segment, and Ile96 and Val122 on the loop regions were colored in magenta, beige, yellow, and green, respectively.
